# Basal ganglia and cortical control of thalamic rebound spikes

**DOI:** 10.1101/386920

**Authors:** Mohammadreza Mohagheghi Nejad, Stefan Rotter, Robert Schmidt

## Abstract

Movement-related decreases in firing rate have been observed in basal ganglia output neurons. They may transmit motor signals to the thalamus, but the effect of these firing rate decreases on downstream neurons in the motor thalamus is not known. One possibility is that they lead to thalamic post-inhibitory rebound spikes. However, it has also been argued that the physiological conditions permitting rebound spiking are pathological, and primarily present in Parkinson’s disease. As in Parkinson’s disease neural activity becomes pathologically correlated, we investigated the impact of correlations in basal ganglia output on the transmission of motor signals using a Hodgkin-Huxley model of thalamocortical neurons. We found that such correlations disrupt the transmission of motor signals via rebound spikes by decreasing the signal-to-noise ratio and increasing the trial-to-trial variability. We further examined the role of sensory responses in basal ganglia output neurons and the effect of cortical excitation of motor thalamus in modulating rebound spiking. Interestingly, both could either promote or suppress the generation of rebound spikes depending on their timing relative to the motor signal. Finally, we determined parameter regimes, such as levels of excitation, under which rebound spiking is feasible in the model, and confirmed that the conditions for rebound spiking are primarily given in pathological regimes. However, we also identified specific conditions in the model that would allow rebound spiking to occur in healthy animals in a small subset of thalamic neurons. Overall, our model provides novel insights into differences between normal and pathological transmission of motor signals.

## Introduction

The basal ganglia have long been implicated in the selection and execution of voluntary movements (Albin et al., 1989; Alexander and Crutcher, 1990b; Redgrave et al., 1999; Hikosaka et al., 2000). Classic “box-and-arrow” models of the basal ganglia (Alexander and Crutcher, 1990a; Wichmann and DeLong, 1996) presume a propagation of motor signals through the direct pathway. The direct pathway consists of direct, inhibitory projections from the striatum to the basal ganglia output regions. Therefore increased activity in the striatum reduces the activity e.g. in the substantia nigra pars reticulata (SNr) (Kravitz et al., 2010). SNr in turn disinhibits the motor thalamus (Deniau and Chevalier, 1985), and thereby enables movement. Basal ganglia output neurons often have high baseline firing rates and decrease their rate during movement in both rodents and primates (Hikosaka and Wurtz, 1983; Schultz, 1986; Leblois et al., 2007; Schmidt et al., 2013). However, recent studies have suggested a more complex picture on how basal ganglia output affects motor thalamus and motor cortex (Bosch-Bouju et al., 2013; Goldberg et al., 2013).

Three different modes have been proposed for how the basal ganglia output can affect thalamic targets (Goldberg et al., 2013). In the first mode sudden pauses in basal ganglia inhibition of thalamus could lead to “rebound” spikes in thalamocortical neurons due to their intrinsic T-type Ca^2+^ channels (Llinás and Jahnsen, 1982). T-type Ca^2+^ channels are de-inactivated during long-lasting hyperpolarization (e.g. due to inhibition from the basal ganglia output), and then activated during the release of this hyperpolarisation (e.g. during a pause in basal ganglia output), which depolarises the membrane potential of thalamocortical neurons. For strong enough preceding hyperpolarisation, the membrane potential can even reach the spike threshold without any excitation (Person and Perkel, 2005; Person and Perkel, 2007; Leblois et al., 2009; Kim et al., 2017). However, thalamocortical neurons also receive excitatory input from cortex, which can change the effect of nigrothalamic inhibition. For moderate levels of cortical excitation the nigrothalamic transmission could operate in a disinhibition mode, in which the basal ganglia effectively gate cortical excitation, so that during pauses of inhibition the excitatory inputs can evoke spikes in the thalamocortical neuron (Kojima and Doupe, 2009; Bosch-Bouju et al., 2014; Edgerton and Jaeger, 2014). If the cortical excitation is strong enough, it is possible that inhibition from the basal ganglia can no longer prevent action potentials in the thalamocortical neurons, but instead controls their timing. In this “entrainment” mode the thalamocortical neuron spikes after the inhibitory input spikes from SNr with a short, fixed latency (Goldberg and Fee, 2012; Goldberg et al., 2012).

One prominent feature of the basal ganglia network is that neurons fire in an uncorrelated fashion, despite their overlapping dendritic fields and local recurrent connections (Wilson, 2013). Specific features of the basal ganglia such as pacemaking neurons and high firing rate heterogeneity may act as mechanisms for active decorrelation of activity. This effectively prevents correlations among neurons, and disrupting this mechanism leads to pathologically correlated activity as in Parkinson’s disease (Bar-Gad et al., 2003; Wilson, 2013). Increased correlated activity has also been observed in basal ganglia output neurons in Parkinson’s disease (Bergman et al., 1998), which can in turn increase correlated activity in the thalamus (Reitsma et al., 2011). Previous computational modelling has shown that pathological basal ganglia output can prevent the thalamic relaying of cortical excitatory signals (Guo et al., 2008). Here we examined how pathological correlations in the basal ganglia output affect the transmission from the basal ganglia to the thalamus, and how this transmission is affected by cortical excitation. We assumed that the movement-related decreases in basal ganglia output transmit a motor signal to the thalamus and that this transmission operates in the rebound transmission mode. In addition to transmitting motor signals, basal ganglia output neurons may also be involved in further sensory and cognitive processing. For example, SNr neurons also respond to salient sensory stimuli instructing the initiation or stopping of movements (Pan et al., 2013; Schmidt et al., 2013). Therefore, we also investigated how these sensory responses may affect the motor transmission.

In the present study we used computational modelling to study the transmission from the basal ganglia to the thalamus via postinhibitory rebound spikes, with a focus on situations in which the basal ganglia output activity exhibits sudden movement-related pauses in activity. We found that for uncorrelated basal ganglia output this transmission had a high fidelity with low trial-to-trial variability in the thalamic response latency, but also occurred only under specific conditions with respect to synaptic connectivity, strength, and firing patterns. In contrast, pathological correlations in SNr strongly increased thalamic rebound spiking and led to a noisy transmission with high trial-to-trial variability. In addition, we found that sensory responses in SNr can, depending on their timing relative to the movement-related decrease, either facilitate or suppress rebound spikes. Therefore, in situations in which rebound spikes play a role for the transmission of motor signals, uncorrelated activity and sensory responses in the basal ganglia output would support the coordinated transmission of motor signals. Finally, we found that the rebound spiking mode persisted in the presence of excitation that is strong enough to maintain baseline firing rates reported in vivo (Bosch-Bouju et al., 2014), and we discuss implications for rebound spiking under healthy and pathological conditions.

## Materials and Methods

### Model neuron

In this study we used a Hodgkin-Huxley type model of a thalamocortical neuron (Rubin and Terman, 2004). The model has four different ionic currents: a leak current (*I*_*L*_), a Na^+^ current (*I*_*Na*_), a K^+^ current (*I*_*K*_), and a T-type Ca^2+^ current (*I*_*T*_), which are determined by the membrane potential *v*, the channel conductances *g* and reversal potentials *E*). While the conductance of the leak current *g*_*L*_ is constant, the conductance of the Na^+^, K^+^ and T-type Ca^2+^ currents depends on the membrane potential and varies over time. These voltage-dependent conductances are formed by the product of the maximum channel conductance (*g*_*Na*_, *g*_*K*_ and *g*_*T*_) and the voltage-dependent (in)activation variables (*m, h, p* and *r*).

The model neuron’s membrane potential is described by

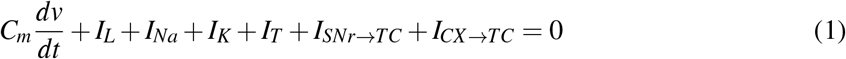

with a leak current *I*_*L*_ = *g*_*L*_[*v*−*E*_*L*_]. The Na^+^ current 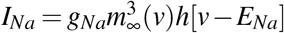 has an instantaneous activation gating variable 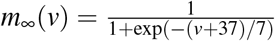 and a slow inactivation gating variable *h* with 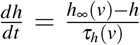 and steady-state 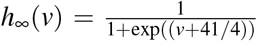 that is approached with a time constant 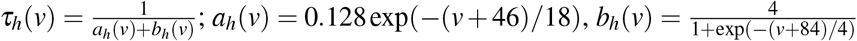.

The activation variable of the K^+^ current *I*_*K*_ = *g*_*K*_ [0.75(1 − *h*)^4^][*v* − *E*_*K*_] is described in analogy to the Na^+^ inactivation variable (*h*), which reduces the dimensionality of the model by one differential equation (Rinzel, 1985a).

The T-type Ca^2+^ current 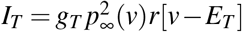 has an instantaneous activation 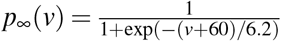 and slow inactivation 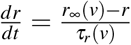 with the steady-state 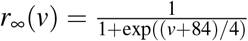 and time constant *τ*_*r*_(*v*) = 28 + 0.3(−(*v* + 25)/10.5). Accordingly, these steady states reach half their maximum at -84mV (*p*_∞_) and -60mV (*r*_∞_), respectively. In Destexhe et al. (1998) slightly different values (−80mV instead of -84mV and -56mV instead of -60mV) were used. Using these values in our model did not lead to substantial changes in rebound spiking behavior of our model (Supplemental Figure 1).

**Figure 1.**
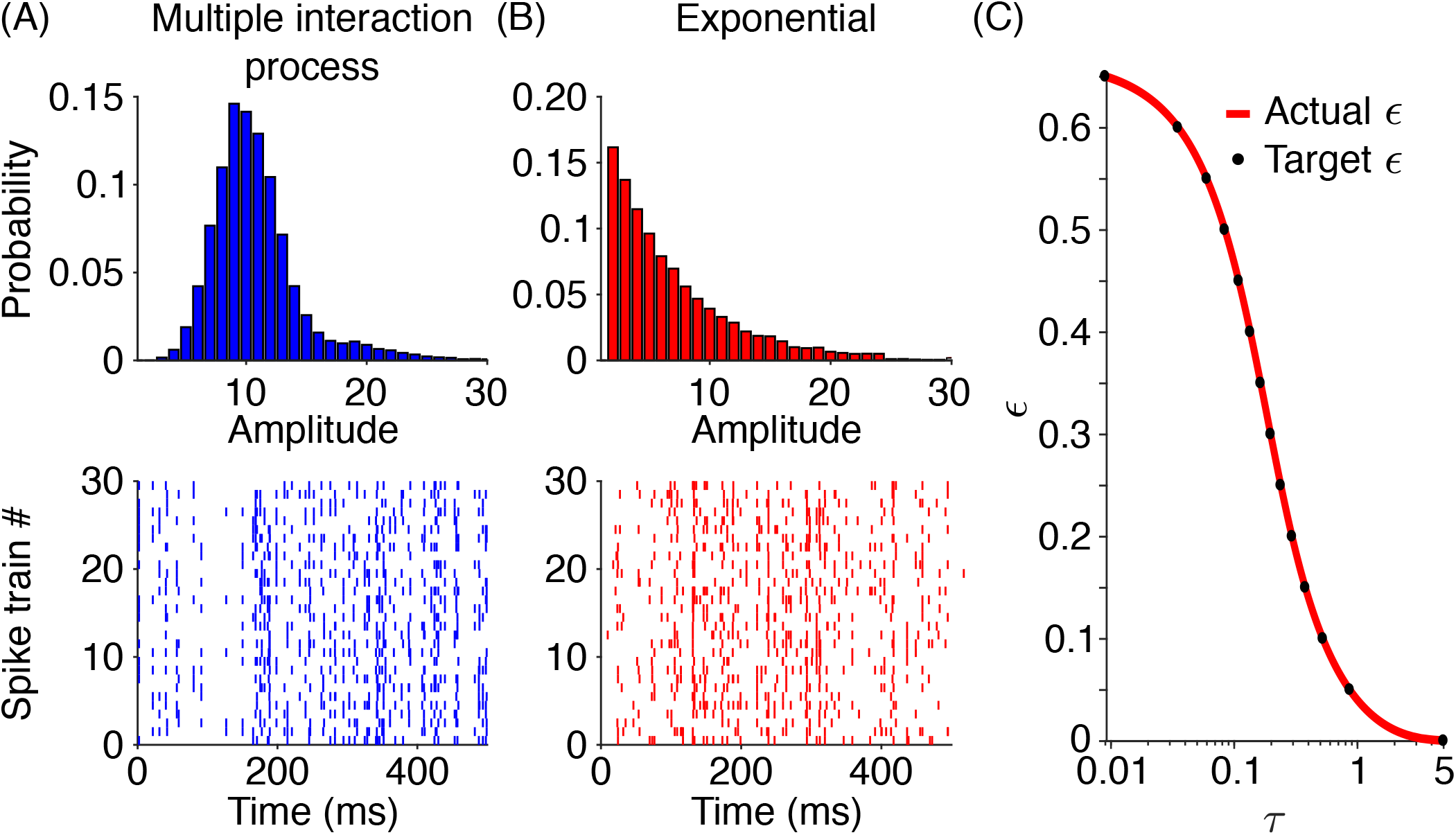
Generation of correlated Poisson spike trains used as input to the model neuron. (A, top) The event amplitude distribution of the higher-order correlations was determined for spike trains generated by a multiple interaction process with *ε* = 0.3 and *r* = 50 Hz. The bottom panel shows the raster plot of 30 respective example spike trains. (B, top) Alternatively, the event amplitude distribution of higher-order correlations followed an exponential amplitude distribution with *ε* = 0.3 and *r* = 50 Hz, and corresponding example spike trains (bottom panel). (C) The parameter *τ* of the exponential amplitude distributions determined the resulting average pairwise correlation *ε* (red trace). Black dots represent the average pairwise correlations that we used to generate input spike trains with an exponential amplitude distribution.

The T-type Ca^2+^ channel can cause post-inhibitory rebound spikes by the following mechanism. Prolonged hyperpolarisation leads to de-inactivation of the T-type Ca^2+^ channel, i.e. the inactivation gate (*r*) opens while the activation gate (*p*) closes. After shutting down the hyperpolarisation, the inactivation gate closes slowly whereas the activation gate opens very fast. Therefore, while both gates are open, the T-type Ca^2+^ conductance increases, inducing an inward current (described by *I*_*T*_) that leads to membrane depolarisation. If this depolarisation is strong enough, this can lead to Na^+^ spikes, which are then referred to as post-inhibitory rebound spikes.

The thalamic model neuron receives two types of synaptic inputs; one inhibitory from the basal ganglia output region SNr (*SNr* → *TC*) and one excitatory from cortex (*CX* → *TC*). Synaptic currents *I*_*X*_ are described by a simple exponential decay with the decay rate *β*_*X*_, where *X* denotes the synapse type (Gerstner and Kistler, 2002). Similar to the intrinsic ionic currents, each synaptic current is described in terms of the membrane potential *v*, channel conductance *g*_*X*_, and the reversal potential *v*_*X*_ : *I*_*X*_ = *g*_*X*_ [*v* − *v*_*X*_] ∑ _*j*_ *s* _*j*_ ; *X* = *{SNr* → *TC,CX* → *TC}*. When a presynaptic neuron *j* spikes at time *t*_*i*_, *s*_*j*_ becomes 1 and decays with time constant *β* afterwards 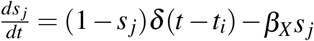, where δ(*t*) is the Dirac delta function. With the conductance caused by a single presynaptic spike (*s*_*j*_ = 1) given by *g*_*X*_, the net synaptic current is therefore the sum of all presynaptic events *s*_*j*_ multiplied by *g*_*X*_ and the difference between the membrane potential and synaptic reversal potential. In our model, the reversal potential for the inhibitory synapse is *v*_*SNr*→*TC*_ = −85*mV* (Rubin and Terman, 2004). With the given parameter settings, our model can evoke rebound spikes for *v*_*SNr*→*TC*_ ≤ −81*mV* (Supplemental Figure 1B). Therefore our model assumes a very low GABA reversal potential, which is, however, in the range of experimentally measured reversal potentials in thalamocortical neurons in the thalamus (Huguenard and Prince, 1994; Ulrich and Huguenard, 1997; Herd et al., 2013). Since there is still some uncertainty on the GABA reversal potential in different thalamic neurons, we checked whether the inhibitory postsynaptic potential evoked by such hyperpolarised reversal potentials is similar to the inhibitory postsynaptic potentials recorded from rats’ motor thalamus in vitro. For the parameter settings used in our study (here 30 synchronous spikes and a GABA maximum conductance of 1 *nS*/*µm*^2^), we found that the inputs in this synaptic settings hyperpolarised the membrane by -17mV, which is very similar to in vitro recordings from rats’ motor thalamus (−18mV, Edgerton and Jaeger, 2014; their Figure 5B). Thalamic rebound spikes can also be driven by the basal ganglia in vivo (Kim et al., 2017), which is in line with very low GABA reversal potentials enabling rebound spikes. The intrinsic and synaptic parameters of the model neuron are described in Table 1.

**Table 1.** Model parameters

| Parameter type | Parameter, value and unit |
| --- | --- |
| Ionic channel conductance | $g_L = 0.05 \text{ nS}/\mu\text{m}^2$<br>$g_{Na} = 3 \text{ nS}/\mu\text{m}^2$<br>$g_T = 5 \text{ nS}/\mu\text{m}^2$<br>$g_K = 5 \text{ nS}/\mu\text{m}^2$ |
| Ionic channel reversal potential | $E_L = -70 \text{ mV}$<br>$E_{Na} = 50 \text{ mV}$<br>$E_T = 0 \text{ mV}$<br>$E_K = -90 \text{ mV}$ |
| Synaptic reversal potential | $v_{SNr \rightarrow TC} = -85 \text{ mV}$<br>$v_{CX \rightarrow TC} = 0 \text{ mV}$ |
| Synaptic decay constant | $\beta_{SNr \rightarrow TC} = 0.08 \text{ ms}^{-1}$<br>$\beta_{CX \rightarrow TC} = 0.18 \text{ ms}^{-1}$ |

Parameters were taken from Rubin and Terman, 2004 and Ermentrout and Terman, 2010.

### Input spike trains

We generated uncorrelated and correlated Poisson spike trains as inputs to the model neuron. To generate uncorrelated spike trains we simulated *N* independent Poisson processes, each with a firing rate *r*. To generate correlated spike trains with a given average pairwise correlation (denoted by *ε*), we considered that for more than 2 input spike trains (*N* ≥ 3), different realisations of spike trains with different correlations of order 3 or higher are possible (Kuhn et al., 2003). For a convenient parametrisation of the order of correlation in input spike trains, we used the distribution of the number of coincident spikes in a time bin, referred to as “event amplitudes” (*A*) (Staude et al., 2010). For a homogeneous population of Poisson spike trains, the average pairwise correlation depends on the first two moments of the amplitude distribution *f*_*A*_:

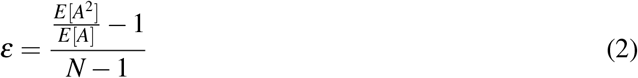

In the present study, we considered binomial and exponential amplitude distributions (Figure 1). While the binomial amplitude distribution has a high probability density around the mean of the distribution (Figure 1A), the exponential distribution has a higher probability density toward smaller amplitudes (Bujan et al., 2015, Figure 1B).

To generate spike trains with a binomial amplitude distribution we implemented a multiple interaction process (Kuhn et al., 2003, Figure 1A). For correlated outputs (*ε* > 0), this was done by first generating a so-called “mother” spike train, a Poisson spike train with rate *λ*. We then subsampled from this mother spike train to derive the set of spike trains used in our simulations as convergent inputs to the model neuron. Each spike train in this set was derived by randomly and independently copying spikes of the “mother” spike train with probability *ε*. The firing rate of each spike train generated via this algorithm is *r* = *ελ*.

We also generated spike trains using exponentially distributed amplitudes described by:

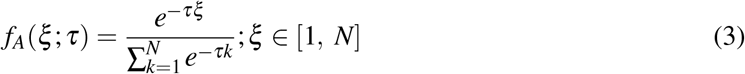

where *f*_*A*_(*ξ* ; *τ*) is the probability density function of event amplitudes *ξ* with the decay rate parameter *τ*. According to Eq. 2, to compute *ε* for this distribution, we needed to compute the proportion of the second moment to the first moment for this distribution. We used the moment-generating function 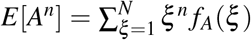 to compute the first and second moments of the distribution and then applied it into Eq. 2, rewriting it to

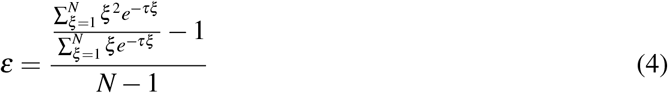

This equation shows that *ε* depends on *τ* and we took a simple numerical approach to find *τ* for each desired *ε*. We computed *ε* for a range of *τ* (from 0 to 5 with steps of 0.001) and then selected the *τ* that yielded an *ε* closest to our desired *ε* (Figure 1C). The maximum error between the *ε* we calculated using Eq. 4 and the desired *ε* was 5 × 10^−4^.

The next step was to generate the population spike trains using the probability distribution determined by the *τ* we already computed. We drew *N* independent Poisson spike trains each for a given event amplitude *ξ* with rate *r*_*ξ*_ = *Nr f*_*A*_(*ξ*)*/ξ* ; *ξ* ∈ [1, *N*]. Since *ξ* represents the number of coincident spikes in a time bin, spike times from independent spike trains should be copied *ξ* times to get the final population spike train used as inputs to the model neuron. As the amplitude distribution described in Eq. 3 has a high probability density toward lower amplitudes, high average pairwise correlations cannot be achieved. For typical parameters of the inhibitory input spike trains in this study (*N* = 30, *r* = 50 Hz), the maximum average pairwise correlation was less than 0.65 (Figure 1C).

### Input spike trains with mixture of binomial and exponential amplitude distributions

We computed the event amplitude distribution of SNr model neurons using a large-scale network model of the basal ganglia (Figure 2D; see also below). This amplitude distribution involved a mixture of exponential and binomial distributions leading to an average pairwise correlation of 0.6 (black dot in Figure 2). To obtain spike trains following this mixed distribution, we first created one spike train with an exponential amplitude distribution contributing 20% of the spikes with an average pairwise correlation of 0.25. Next, another spike train with a binomial amplitude distribution was generated (see above), contributing the remaining 80% of the spikes in the input spike train. We changed the average pairwise correlations of these input spike trains by only changing the average pairwise correlation of the subset with the binomial amplitude distribution.

**Figure 2.**
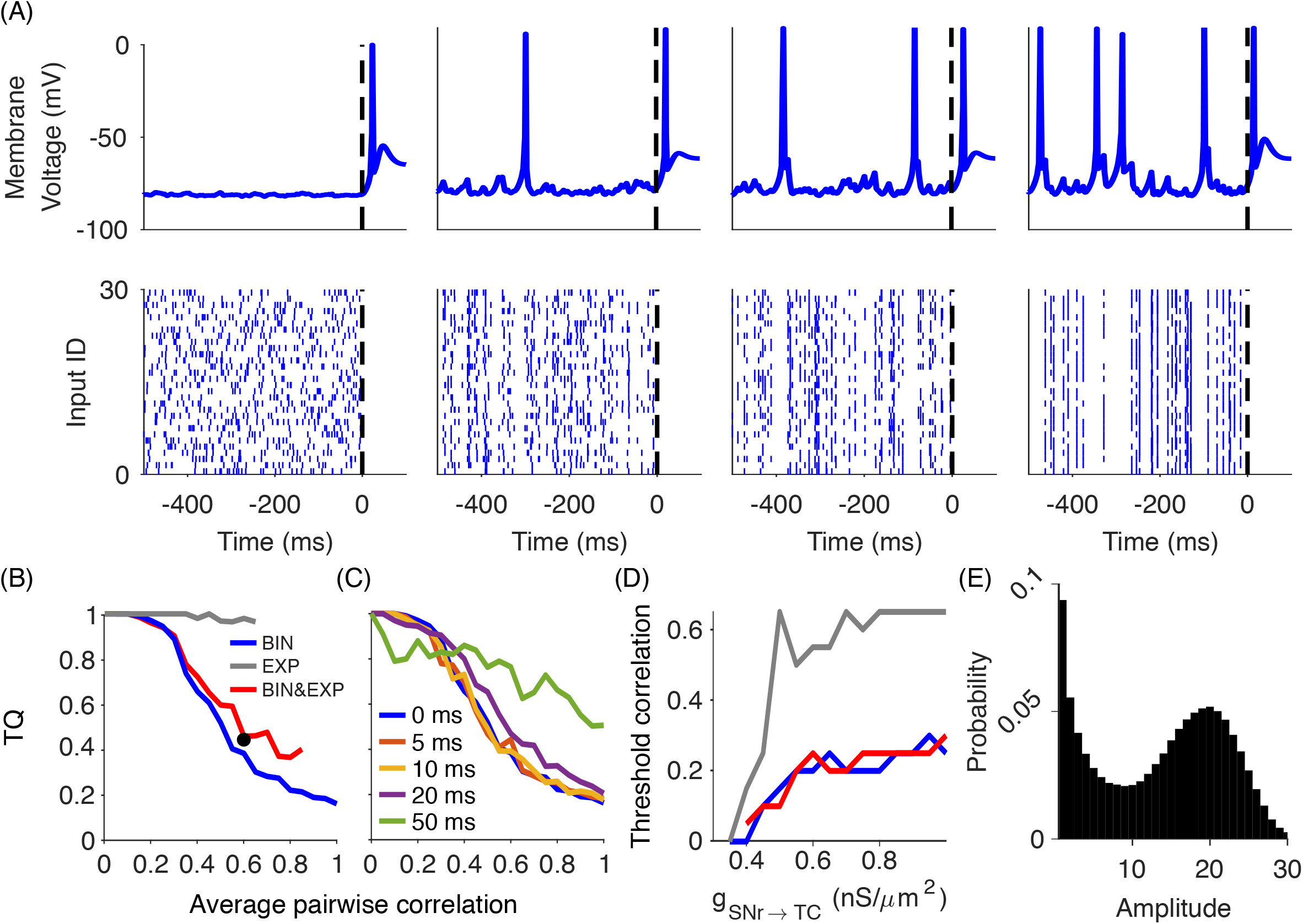
Input spike correlations impair the transmission quality (TQ) of motor signals from SNr to thalamus. (A) Top panels show the intracellular response of the thalamocortical model neuron to the inhibitory input spike trains from SNr displayed in the bottom panels. Uncorrelated Poisson spike trains (*ε* = 0) led to high-fidelity transmission (TQ = 1) via a single rebound spike after the firing rate decrease in the input (leftmost panel). Correlated Poisson spike trains, however, led to rebound spikes at random times, whenever there is a pause in the input spike trains (left middle panel: *ε* = 0.2 leading to TQ = 0.5, right middle panel: *ε* = 0.35 leading to TQ = 0.33 and rightmost panel: *ε* = 0.7 leading to TQ = 0.25). (B) Impact of input correlations on TQ depended on the correlation model (BIN, binomial; EXP, exponential; BIN&EXP, mixture of both). Note that the exponential distribution of the event amplitudes had a maximum average pairwise correlation of 0.65 (see Materials and Methods). The black dot marks the TQ for the spike trains generated using the event amplitude distribution shown in (E). (C) For the binomial correlation model, jittering the input spike times decreased the TQ only for long jitter time windows (50ms), indicating that correlations on longer time scales are overall less detrimental. (D) The threshold correlation at which the transmission quality deteriorated (TQ*<* 0.95) only weakly depended on the inhibitory input strength (same legend as in B). (E) The simulation of Parkinson’s disease in a large-scale model of the basal ganglia yielded an event amplitude distribution of SNr spike times that corresponded to a mixture of the exponential and binomial amplitude distributions.

### Uncorrelated input spike trains with gradual decrease

We captured the gradual movement-related decrease, which is observed experimentally, by using a sigmoid function to describe the firing rate of the input spike trains as a function of time 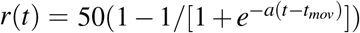 Hz. We varied the slope parameter, *a*, to change the slope of the firing rate decrease. *t*_*mov*_ is the time point (in this study at one second), when the firing rate decreases to the half maximum, i.e. *r*(*t*_*mov*_) = 25 Hz.

### Data analysis: identifying rebound spikes

The model neuron can fire spikes in response to excitatory input or due to release from inhibition with post-inhibitory rebound spikes. Therefore, one challenge was to distinguish “normal” spikes driven by excitatory inputs from post-inhibitory rebound spikes. In mice studies, genetic approaches are often used to knockout T-type Ca^2+^ channels, which are critical for generation of post-inhibitory rebound spikes (Kim et al., 2017). We adopted this in our model by simply removing the T-type Ca^2+^ channels in our model (i.e. *g*_*T*_ = 0 *nS/µm*^2^). However, this also caused changes in the intrinsic properties of the model neuron such as its excitability. We therefore took a more elaborate approach tailored to each of the two excitation scenarios, single excitatory spikes (Figure 5) and spontaneous excitation (Figure 6).

For the simulations with a single excitatory input spike the identification of rebound spikes was straightforward because the used excitatory strengths were subthreshold and thus could evoke no spikes. Therefore, we labelled all generated spikes as rebound spikes. However, for the simulations with ongoing excitation, the excitatory input was able to evoke “normal” spikes as well. To identify rebound spikes there, we simulated the model neuron with three different input combinations, inhibition-only, excitation-only and inhibition-excitation. For inhibition-only input, we determined the output firing rate of the model neuron purely due to rebound spiking (*f*_*I*_). In addition, we determined the time window in which the model neuron fired those rebound spikes (as this was typically in a short time window just after the movement-related decrease). We then compared the rebound-driven firing rate in this time window with the firing rate *f*_*E*_ obtained from an excitation-only simulation (i.e. without any inhibitory input, so no rebound spikes). Finally, we fed our model with both inputs (inhibition-excitation) and computed the firing rate in that time window, which involved both rebound and non-rebound spiking (*f*_*EI*_). We then computed the proportion of rebound spiking as: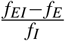.

### Data analysis: transmission quality

For our simulations shown in Figure 2, we needed to quantify the transmission quality for a variety of inputs strengths and degrees of correlation. For a high-fidelity transmission of the motor signal the thalamocortical neuron would ideally respond only to the movement-related decrease of activity in SNr neurons with a rebound spike, and be silent otherwise. Any rebound spike before the movement-related decrease would make the transmission noisy, in the sense that the decoding of the presence and timing of the motor signal in thalamic activity would be less accurate. Therefore, we used the number of spikes after the onset of the movement-related decrease, normalised by the total number of spikes within -1 s to 0.5 s around the onset of the movement-related decrease as a measure of the transmission quality.

### Large-scale model of the basal ganglia

We utilised a large-scale network model of the basal ganglia (Lindahl and Kotaleski, 2016) to compute the distribution of event amplitudes in SNr during pathological activity in dopamine-depleted basal ganglia. This network model mimics the pathological activity pattern observed experimentally in a rat model of Parkinson’s disease. To achieve the pathological activity pattern in SNr, we ran this model using a default parameter set originally from this network model. This parameter set involved setting dopamine modulation factor to zero and inducing a 20-Hz modulation to the emulated cortical inputs to the basal ganglia regions (for details see Lindahl and Kotaleski, 2016).

### Software packages

We implemented the model neuron in Simulink, a simulation package in MATLAB (R2016b) and used a 4th-order Runge-Kutta method to numerically solve the differential equations (time step = 0.01 ms). We wrote all scripts to generate input spike trains, handle simulations and analyse and visualise the simulation data in MATLAB. For our simulations we used the bwForCluster NEMO, a high-performance compute resource at Freiburg University.

## Results

### Uncorrelated activity prevents rebound spiking

Correlated activity in the basal ganglia is usually considered pathological (Bergman et al., 1998; Bar-Gad et al., 2003; Wilson, 2013) and might lead to the generation of rebound spikes (Edgerton and Jaeger, 2014). To determine how correlated activity in basal ganglia output affects rebound spiking, we simulated a thalamocortical neuron exposed to inhibitory Poisson input spike trains with varying degrees of correlation (Figure 2). Including only inhibitory inputs, in the first step, enabled us to elucidate the characteristics of inhibition that are essential for generating rebound spikes and facilitated the identification of post-inhibitory rebound spikes by excluding the possibility for evoking excitatory-driven spikes. We used binomial and exponential amplitude distributions to generate correlated Poisson spike trains (see Materials and Methods). In addition, we modulated the input firing rate so that it mimicked the prominent movement-related decrease of basal ganglia output neurons observed in experimental studies (Hikosaka and Wurtz, 1983; Schultz, 1986; Leblois et al., 2007; Schmidt et al., 2013).

For uncorrelated inputs the model responded to the movement-related decrease with a single rebound spike (Figure 2A, left panel). However, for correlated inputs rebound spikes appeared not only after the movement-related decrease, but also at random times during baseline activity (Figure 2A, middle and right panels). The reason for this was that correlated SNr activity led not only to epochs with many synchronous spikes, but also to pauses in the population activity that were long enough to trigger rebound spikes.

While studies on songbirds suggest strong one-to-one projections from Area X (basal ganglia output equivalent in avians) to the medial portion of the dorsolateral nucleus of the anterior thalamus (DLM) (Person and Perkel, 2005; Leblois et al., 2009), in rats multiple inhibitory projections from SNr converge on a single thalamocortical neuron (Edgerton and Jaeger, 2014), which affects the strength of the inhibition on the thalamocortical neuron. Electron microscopic studies of these projections show a similar synaptic structure in rats and monkeys suggesting that the rodent nigrothalamic pathway can be a valid model for studying GABAergic transmissions in primates (Bodor et al., 2008). Similar to the synaptic strength, the precise degree of nigrothalamic convergence is not known. Edgerton and Jaeger (2014) estimated that 3-13 SNr neurons project to a single thalamocortical neuron. As they noted that this may be an underestimate due to experimental limitations of opsin expression, we chose a larger number (30) for the degree of convergence. However, in our model rebound spikes also occurred for a smaller number of inputs (20<N<30; Supplemental Figure 2). To determine whether these factors are relevant for our findings on the transmission quality, we repeated our simulations for different inhibitory strengths, but found that the transmission quality did not depend on the inhibitory strength as long as the inhibition was strong enough to lead to rebound spikes (Figure 2D). As for more than two inputs the input spike trains cannot be uniquely characterised by pairwise correlations, we considered two different possibilities for higher-order correlations (see Materials and Methods). We found that the transmission quality strongly depended on both the input average pairwise correlation and higher-order correlations among input spike trains (Figure 2B).

Pairwise correlations affected the transmission for a binomial amplitude distribution (Figure 2B, dark blue trace). For a binomial amplitude distribution higher-order events (“population bursts”) are common, which increases the probability for pauses in the population activity. Thereby, even weak correlations among SNr spike trains led to a sharp decrease in the transmission quality. In contrast, for spike train correlations with an exponential amplitude distribution, the decrease in transmission quality was less pronounced (Figure 2B, grey trace). This was because for the exponential amplitude distribution lower-order events are more common, which are not sufficient for pauses in the population activity of SNr neurons leading to thalamic rebound spikes. Therefore, in particular higher-order correlations may be responsible for pathological increases in rebound spiking and disrupt motor signalling.

We further investigated whether the substantial decrease in the transmission quality observed for the binomial amplitude distribution depended on millisecond synchrony of correlated spike times. We jittered the synchronous spike events using different time windows (Figure 2C), which corresponds to correlations on slower timescales. We found that the transmission quality decreased for jittering timescales ≤ 20 ms similar to inputs with correlations on a millisecond timescale (i.e. without jittering), confirming that the decrease in transmission quality does not depend on millisecond synchrony. However, correlations on the timescale of 50 ms did not substantially influence the transmission quality, as was expected due to the lack of population pauses.

The purpose of our simulation of correlated activity was to mimic basal ganglia output patterns in Parkinson’s disease. However, as the event amplitude distribution of pathologically correlated activity in SNr is currently unknown, we employed a large-scale model of the basal ganglia (Lindahl and Kotaleski, 2016), in which beta oscillations propagate through cortico-basal ganglia circuits (see Materials and Methods). Beta oscillations are widely observed in animals with dopamine-depleted basal ganglia including their output nuclei (Brown et al., 2001; Avila et al., 2010). While beta oscillations can be generated in the pallido-subthalamic loop (Kumar et al., 2011; Mirzaei et al., 2017), here we did not assume a specific mechanism for the generation of correlated activity in Parkinson’s disease, but focussed on the event amplitude distribution in SNr in a simulation of Parkinson’s disease. We found that the amplitude distributions in the dopamine-depleted state of the large-scale model were somewhere in between binomial and exponential (Figure 2E).

To investigate the model with a correlation structure that might be relevant for Parkinson’s disease, we generated input spike trains based on a mixture of binomial and exponential distributions (see Materials and Methods). We then investigated the effect of different average pairwise correlations in this mixed distribution. We found that increasing the average pairwise correlation of the binomial component of the mixed distribution had a similar effect on the transmission quality as in the standard binomial amplitude distribution (Figure 2B, red and blue traces). Furthermore, for the average pairwise correlation found from the large-scale model for Parkinson’s disease the transmission quality was low (Figure 2B, black dot). This suggests that under a correlation structure similar to Parkinson’s disease, even weak correlations in basal ganglia output may impair the transmission of motor signals in the rebound transmission mode. Whether this mechanism could contribute to motor symptoms in Parkinson’s disease, also depends on the structure of excitatory inputs (Magnin et al., 2000; Edgerton and Jaeger, 2014; Kim et al., 2017).

### Uncorrelated activity increases transmission speed and reduces variability

Having demonstrated that correlated activity can increase pathological rebound spiking, we next examined whether correlations can also affect transmission speed and trial-to-trial variability. We assumed here that movement-related decreases in the firing rate of basal ganglia output neurons transmit motor signals to the thalamus via the rebound transmission mode.

To study the effect of input correlations on transmission speed, we used the same scenario as above (Figure 2) and measured the time between the onset of the movement-related decrease and the rebound spike. We found that the transmission speed was fastest for no or weak correlations, and slower for stronger correlations (Figure 3A). Therefore, at least in this simplified scenario without excitation, uncorrelated activity in basal ganglia output regions may also promote the fast transmission of motor signals. To generalise our findings on the transmission speed beyond the scenario using the movement-related decrease, we further examined transmission speed using (rebound) spike-triggered averages of inputs. Instead of simulating a movement-related decrease, we exposed the model neuron to inhibitory inputs with a constant firing rate. To compute the spike-triggered average, we used the peak of each rebound spike as the reference time point to compute the average of the preceding input. Since rebound spikes occurred more often for stronger input correlations, we performed this analysis on inputs having a correlation coefficient of either 0.3 or 1.0. These simulations confirmed that weak input correlations induce faster transmission than strong correlations (Figure 3C).

**Figure 3.**
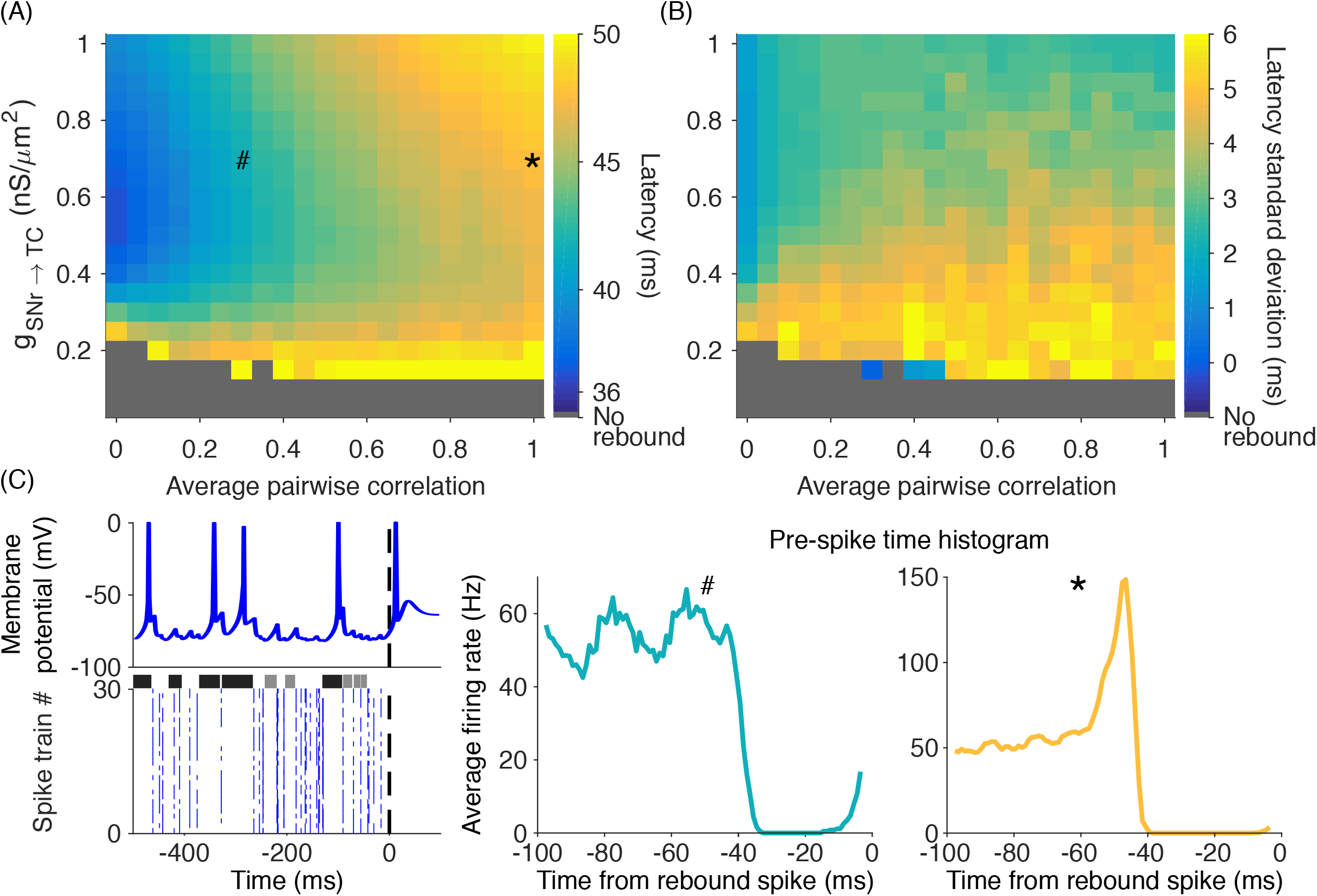
Correlated SNr spike trains decrease transmission speed and temporal precision of rebound spikes. Systematic investigation of average transmission latency (A) and its standard deviation (B) for different degrees of correlation and inhibitory strengths identified the range with fastest transmission speed and highest transmission precision, respectively. (C) Left panel shows a sample membrane potential (*g*_*SNr→TC*_ = 0.70 *nS/µm*^2^, *ε* = 0.7; top) of the thalamocortical model neuron and the corresponding inhibitory inputs (bottom). Note that rebound spikes were preceded by pauses in the input raster plot (indicated by black horizontal bars). However, for very short pauses (indicated by grey horizontal bars) no rebound spikes occurred. Averages triggered by rebound spikes for weakly correlated inputs (C, middle panel) and strongly correlated inputs (C, right panel) confirmed that pauses in the inhibitory input preceded rebound spikes. The duration of the pause preceding the rebound spikes reflected the transmission latency. The inset symbols (#, *) in (A) indicate the parameters used for the corresponding spike-triggered averages in (C).

For the transmission of motor signals via rebound spikes the trial-to-trial variability of the transmission speed may be important. For example, to coordinate motor signals across different neural pathways low variability (i.e. high precision) of the transmission speed might be necessary. To investigate the nigrothalamic transmission variability, we computed the variance over the latencies across 100 trials with movement-related decreases in SNr activity (i.e. the same scenario as in Figure 3A). We found that for uncorrelated inputs transmission was very precise in the sense that the trial-to-trial variability of the response latency was small (Figure 3B). In contrast, even weak correlations led to a high transmission variability due to changes in the amount of hyperpolarisation caused by correlated inputs preceding rebound spikes. This simplified scenario, without excitation, suggests that uncorrelated inhibitory inputs may enable a high precision of the transmission via rebound spikes by reducing the trial-to-trial variability in response latency.

### Sensory responses can promote or suppress rebound spiking

SNr neurons often have short-latency responses to salient sensory stimuli characterised by brief increases in firing rate (Pan et al., 2013). In rats performing a stop-signal task these responses also occurred in neurons that decreased their activity during movement (Schmidt et al., 2013). This included responses to auditory stimuli, which cued the initiation of a movement (Go cue) or the cancellation of an upcoming movement (Stop cue). While in these experiments sensory cues prompted movement, we assume here that similar responses also occur in other behavioural contexts. We examined how brief increases in SNr activity, similar to sensory responses, affect transmission in the rebound spiking mode (Figure 4). The thalamocortical model neuron received inputs similar to the SNr firing patterns recorded in rats during movement initiation (i.e. uncorrelated inputs with high baseline firing rate and a sudden movement-related decrease). To model sensory responses in the SNr neurons, we added a brief increase in firing rate at different time points relative to the movement-related decrease (Figure 4A). We generated the brief increase by adding a single spike in each spike train having the sensory response at the desired time point. This allowed us to observe the effect of the timing of sensory responses on rebound spiking.

**Figure 4.**
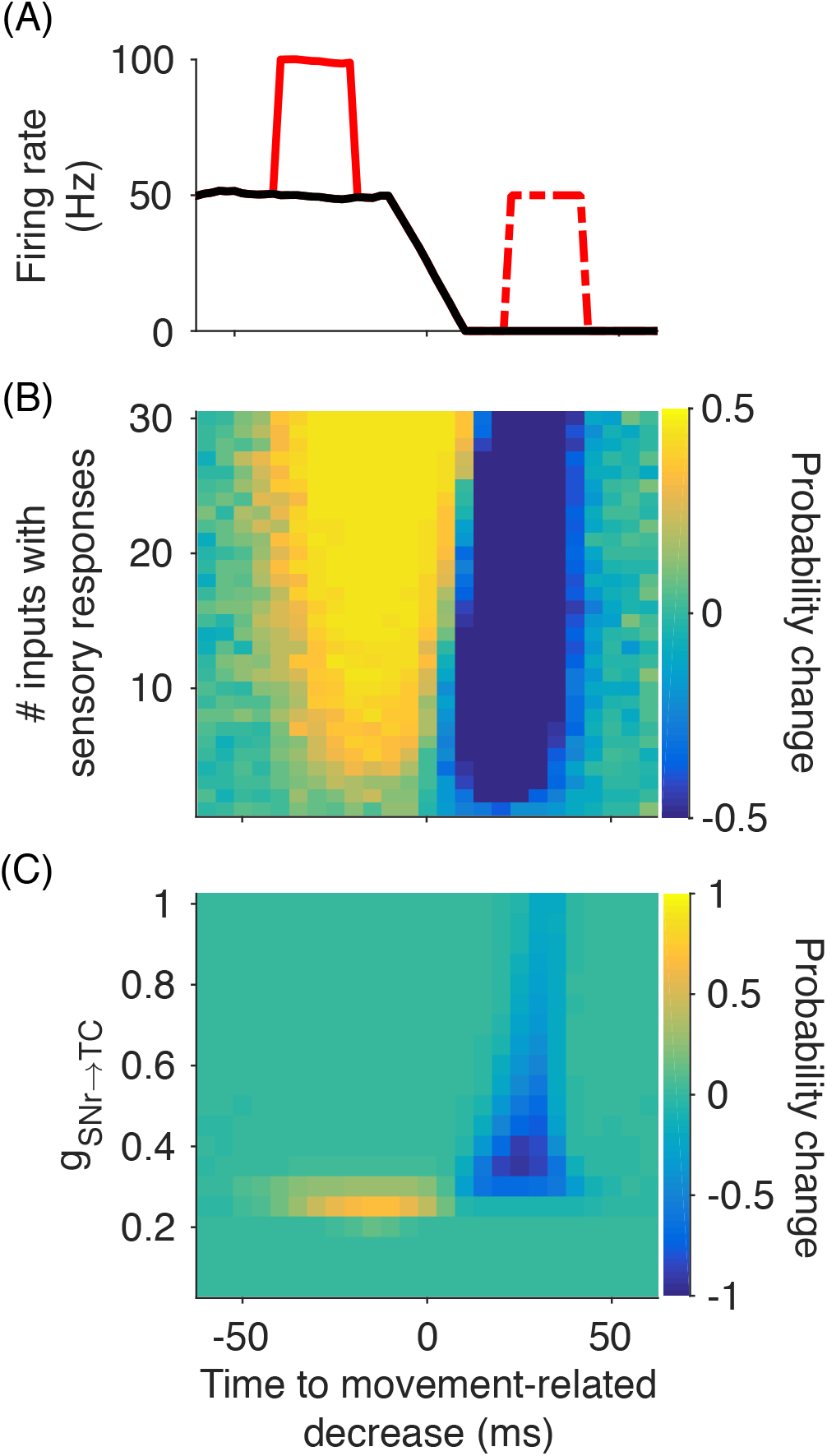
Sensory responses in SNr firing rate change the probability of rebound spikes in the thalamocortical model neuron. (A) The simulations used an average firing rate as input, which reflected the SNr firing rate with a movement-related decrease (black line). Sensory responses (red lines) were then added to the input at different time points relative to the movement-related decrease. Here two example timings are shown, before (solid) and after (dash-dot) the movement-related decrease. (B) The timing of the sensory responses relative to the movement-related decrease was varied systematically (x-axis). For a given relative timing, we determined whether rebound spikes were suppressed (blue area) or facilitated (yellow area; here *g*_*SNr→TC*_ = 0.29 *nS/µ m*^2^). Note the large impact of the timing of the sensory response on the probability of rebound spikes, even if it occurred in only a small subset of neurons. (C) The input strength *g*_*SNr→TC*_ affects the suppression and facilitation of rebound spikes. Here the change in rebound probability was averaged across the number of inputs with sensory responses (across y-axis in B).

To quantify the effect of sensory responses, we measured the difference in the probability of generating a rebound spike after the movement-related decrease in simulations with and without sensory responses. Interestingly, the sensory responses could either increase or decrease the probability of generating a rebound spike, depending on their relative timing to the movement-related decrease (Figure 4B). For sensory responses preceding the movement-related decrease for up to 40 ms, the probability of generating a rebound spike was increased. This was because the sensory response led to additional hyperpolarisation in the thalamocortical neuron, which promoted rebound spiking. In contrast, for sensory responses occurring 10-40 ms after the movement-related decrease, the probability of generating a rebound spike was decreased. This was because the sensory response in that case partly prevented the movement-related pause of SNr firing. Together, this points to the intriguing possibility that sensory responses in SNr can have opposite effects on behaviour (either promoting or suppressing movement), depending on their timing (Figure 4B). This could explain why SNr neurons respond to both Go and Stop cues with a similar increase in firing rate (Schmidt et al., 2013; Mallet et al., 2016), a previously puzzling finding (see Discussion). However, in an *in vivo* situation, there would likely be additional excitatory inputs to both SNr and the thalamus, which would affect whether rebound spikes are generated in this situation.

In addition to the timing of sensory responses relative to the movement-related decrease, also the inhibitory input strength modulated the probability of generating a rebound spike (Figure 4C). For weaker inhibitory inputs (*g*_*SNr*→*TC*_ = 0.25*nS/µm*^2^), the probability of generating a rebound spike was increased because the additional inhibitory inputs contributed to the hyperpolarisation of the thalamocortical neuron. However, for slightly stronger inputs (*g*_*SNr*→*TC*_ ≥ 0.35*nS/µm*^2^), the sensory responses could not further facilitate rebound spiking because the probability of generating a rebound spike was already one. Accordingly, sensory responses were most effective in reducing the probability of generating a rebound spike for medium input strengths (i.e. with a relatively high probability of generating a rebound spike). We found that the most effective strength for suppressing rebound spikes was at *g*_*SNr*→*TC*_ = 0.35*nS/µm*^2^. However, the suppressing effect vanished for *g*_*SNr*→*TC*_ ≥ 0.8*nS/µm*^2^ because for this strength, without any excitatory inputs, the sensory responses themselves caused a hyperpolarization strong enough to trigger a rebound spike (Figure 4C). Therefore, the effect of sensory responses in SNr on motor signals strongly depended on the nigrothalamic connection strength.

### Rebound spikes in the presence of excitation

Having studied basic properties of rebound spiking in the model under somewhat idealised conditions, we next extended the model to account for further conditions relevant in vivo. For example, we have assumed so far that the thalamocortical neuron receives input from SNr neurons that decrease their activity during movement. However, electrophysiological recordings in SNr and other basal ganglia output neurons have also identified neurons that do not decrease their activity during movement (Schmidt et al., 2013). Therefore, we investigated the response of the thalamocortical model neuron in a scenario in which only a fraction of SNr inputs decreased their firing rates, while the remaining neurons did not change their rates (Figure 5). We found that the thalamocortical model neuron elicited a rebound spike with high probability only when a large fraction of input neurons decreased their firing rates to zero (Figure 5A). If we assume random connectivity, this would mean that only a very small percentage of thalamic neurons receives inputs from a sufficient number of nigral neurons with a movement-related decrease to elicit rebound spikes. Therefore, in order for this mechanism to apply to healthy animals, non-random connectivity would be required, so that different nigral neurons with movement-related decreases in firing rate preferentially converge onto the same thalamic target neuron.

**Figure 5.**
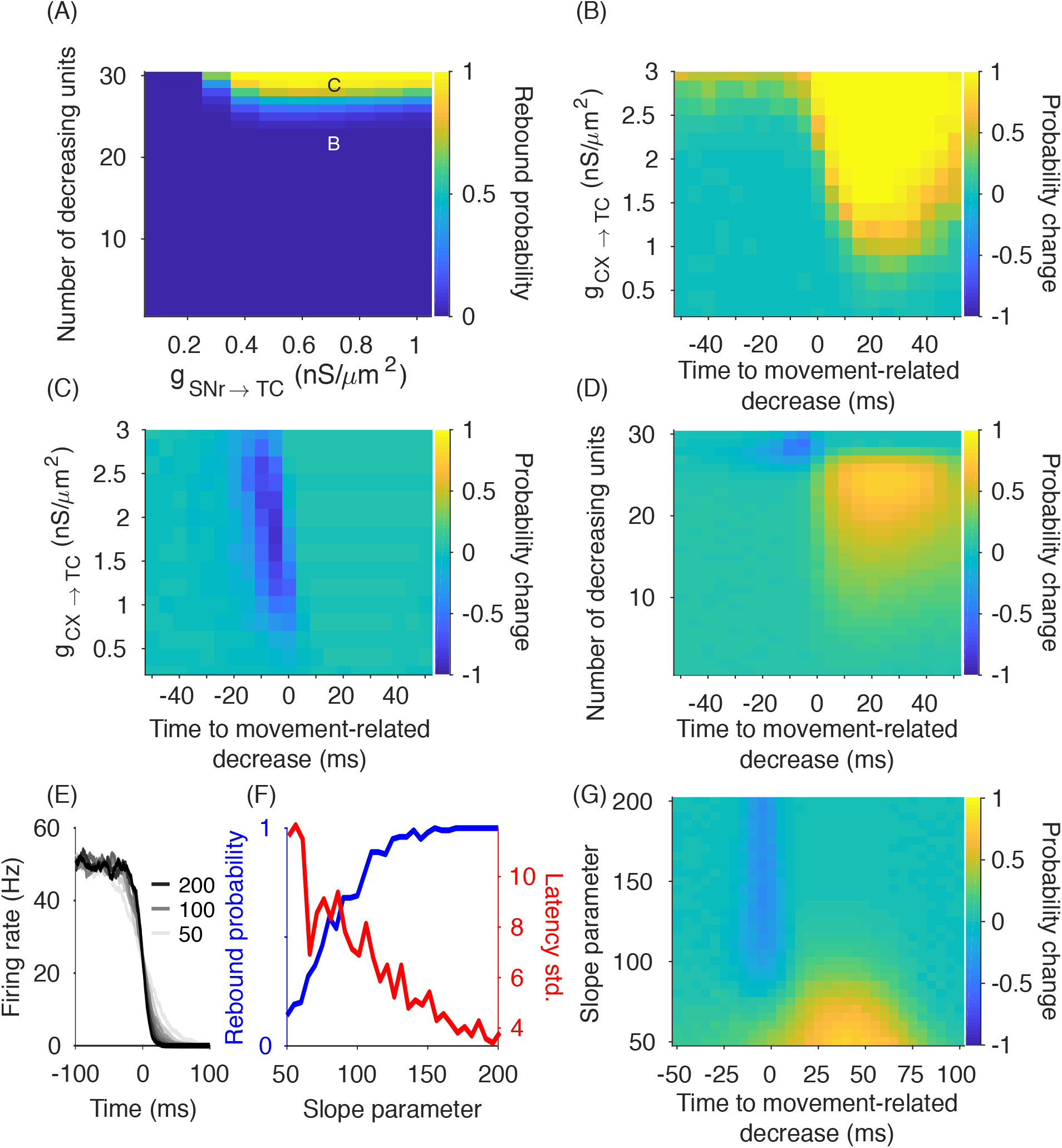
Effect of precisely timed excitatory input spikes on rebound spiking. (A) The generation of rebound spikes requires that a large fraction of the inhibitory input spike trains exhibit a movement-related decrease in firing rate, largely independent of their input strength. (B) Adding a single excitatory spike as input to the thalamocortical model neuron strongly increases the probability of rebound spike generation compared to pure inhibitory inputs (letter “B” in panel A). Note that this occurs in a regime, in which usually no rebound spike can be generated because not enough (here 22 out of 30) neurons decrease their firing rate. (C) In a regime, in which usually rebound spikes are generated (letter “C” in panel A), adding a single excitatory spike as input to the thalamocortical neuron decreases the probability of rebound spike generation compared to pure inhibitory inputs. (D) Systematic investigation of the parameter space indicates a narrow regime, in which a single excitatory spike can decrease, and a larger regime, in which it can increase the probability of a rebound spike. Here, the probability changes are averaged over excitatory input strengths.

The large fraction of SNr neurons required to exhibit a movement-related decrease in order to elicit a rebound spike downstream constrains the scenario under which this transmission is plausible in vivo. However, in a more realistic scenario the thalamocortical neuron also receives excitatory inputs (e.g. from cortex). Therefore, we examined whether excitatory input can, under some conditions, enhance the transmission via rebound spiking (Figure 5B-D). Importantly, the excitatory inputs should be weak enough in order not to elicit spikes themselves. We simulated the model neuron by adding a single excitatory input spike with variable timing with respect to the movement-related decrease in the inhibitory inputs, and observed whether it promoted or suppressed rebound spikes. Using single input excitatory spikes enabled us to accurately determine the minimal excitatory conductance that was required for modulating rebound spikes. While our results below indicate that single spikes can have a powerful effect on modulating rebound spikes, this does not necessarily mean that these processes also rely on single spikes in vivo. We investigated the effect of the excitatory spike on the probability of generating a rebound spike by comparing a simulation including excitatory and inhibitory inputs with a simulation that included only inhibitory inputs. We found that for parameter regions in which the probability of generating a rebound spike was usually small (i.e. in the dark blue region in Figure 5A), additional excitatory spikes after the movement-related decrease increased the rebound probability (Figure 5B). We confirmed that these spikes in the thalamocortical neuron are actually rebound spikes (and not just driven by the excitatory input; see Materials and Methods). However, for strong excitation, the thalamocortical model neuron spiked also before the SNr movement-related decrease, indicating that these spikes were no longer rebound spikes.

For parameter regions in which the probability of generating a rebound spike was high (i.e. outside the dark blue region in Figure 5A), the excitatory input spikes could also suppress the generation of rebound spikes when they occurred before the movement-related decrease (Figure 5C). In contrast, when the excitatory input spike occurred after the movement-related decrease, it enhanced the probability of generating a rebound spike. Therefore, similar to the complex effect of sensory responses in SNr neurons described above, also the excitatory input to the thalamocortical neurons could either promote or prevent rebound spikes depending on its timing. Furthermore, if only a fraction of SNr neurons exhibited a movement-related decrease, precisely timed excitatory input could promote the transmission of motor signals to thalamocortical neurons (Figure 5D). Overall, our simulations indicate that rebound spikes can also occur in a parameter regime that includes excitation. Furthermore, precisely timed excitation provides an additional mechanism of rebound spike modulation. In monkeys performing a learned reaching movement, thalamic excitation seems to precede basal ganglia motor output (Schwab et al., 2020), which according to our model could therefore indicate that excitatory inputs to the thalamus are timed to suppress rebound spiking in healthy animals.

### Role of the slope of the movement-related decrease

So far we assumed that the movement-related decreases in SNr firing rate are abrupt. However, electrophysiological recordings in rodents (Schmidt et al., 2013) and non-human primates (Hikosaka and Wurtz, 1983; Schultz, 1986; Leblois et al., 2007) indicate that, at least in data averaged over trials, the firing rate decreases can also be more gradual. Therefore, we investigated the impact of input spike trains with various slopes (see Methods) on rebound spikes (Figure 5E). We found that steep slopes of the movement-related firing rate decrease led to rebound spikes with high probability and small timing variability (Figure 5F). In contrast, more gradual movement-related decreases reduced the probability of rebound spikes and increased the spike timing variability.

We further investigated the impact of single excitatory spikes (similar to above) on the probability of rebound spikes for different SNr firing rate slopes (Figure 5G). We found that, if the slope was too small to reliably evoke rebound spikes (low rebound probability), excitatory spikes briefly after the onset of the movement-related decrease could increase the probability of rebound spikes. In contrast, for steeper slopes, the probability of rebound spikes decreased when the excitatory spike occurred before the movement-related decrease. These results further support that excitation can powerfully modulate rebound spiking and even promote rebound spikes under circumstances in which the inhibitory input characteristics are by themselves insufficient for the generation of rebound spikes. Furthermore, rebound spiking is reduced, if cortical excitation precedes the movement-related decrease. This may be the case in healthy animals performing learned reaching movements (Schwab et al., 2020).

### Transmission modes revisited: prevalence of rebound spiking

The interaction of excitation and inhibition in thalamocortical neurons is important because even weak excitation may change the transmission mode from rebound to disinhibition (Goldberg et al., 2013). As we observed rebound spiking in the presence of single excitatory spikes (Figure 5), we further investigated how ongoing excitation affects the mode of nigrothalamic transmission. As before, we simulated the model neuron with movement-related inhibitory inputs, but added a background excitation mimicking input from many cortical neurons in the form of a Poisson spike train with the firing rate of 100 Hz and examined the effect of changing excitatory strength (Figure 6). In an idealised scenario the model neuron spikes exclusively after the SNr movement-related decrease for both the rebound and disinhibition transmission modes. These spikes are either post-inhibitory rebound spikes (in the rebound mode), or the result of depolarisation through excitation (in the disinhibition mode). However, we found that rebound and disinhibition modes could also coexist in regimes in which the model neuron has non-zero baseline firing rates (Figure 6A).

**Figure 6.**
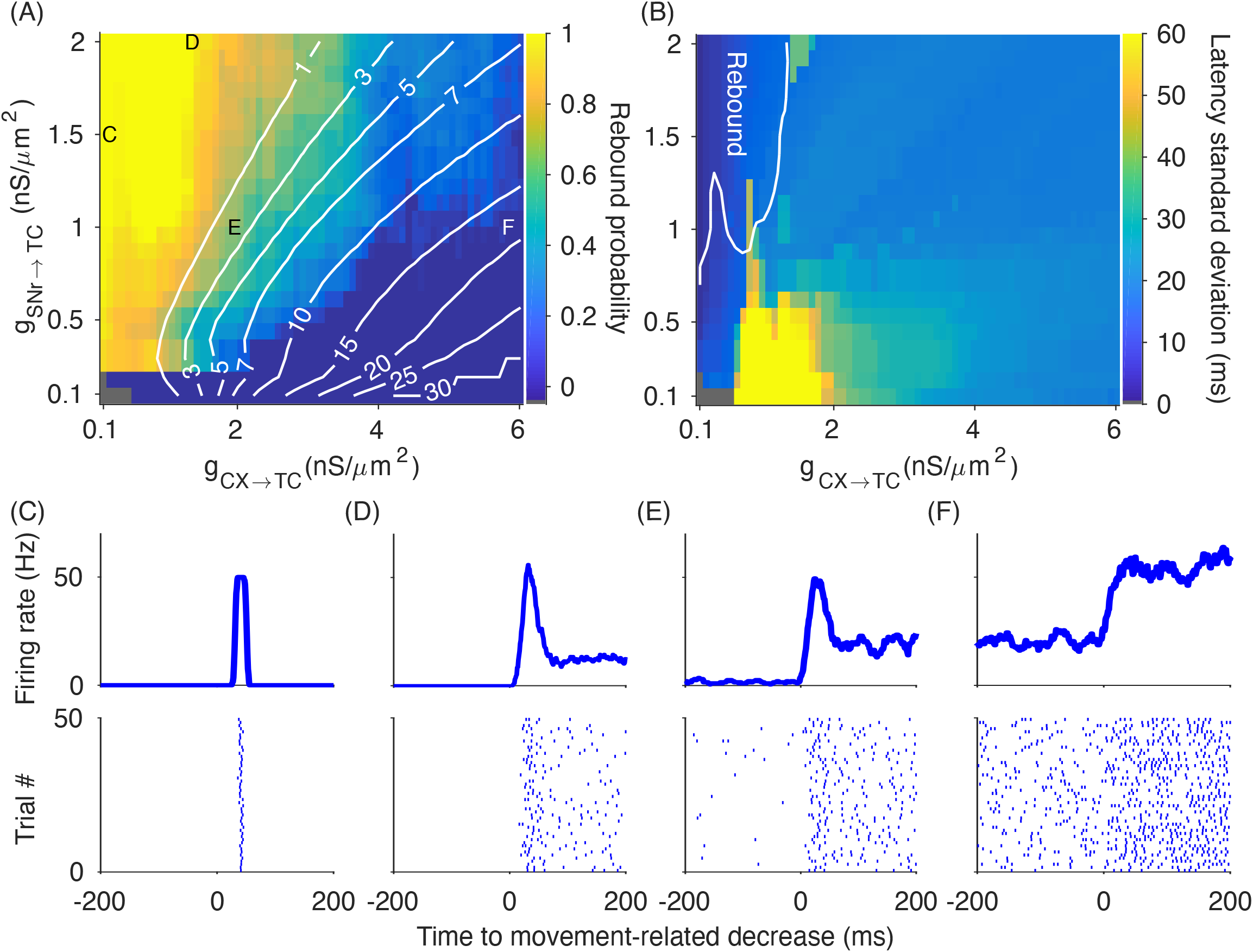
Smooth transition from rebound to disinhibition transmission mode. (A) The probability of rebound spikes only gradually decreased with stronger excitatory inputs, indicating a large parameter regime in which the rebound and disinhibition transmission modes coexisted. The yellow area marks the regime in which transmission was exclusively mediated by rebound spiking, while in dark blue areas the basal ganglia output only disinhibited cortical excitation. The white isolines illustrate the baseline firing rate of the model neuron (i.e. the firing rate before the onset of the movement-related decrease in the input). In the small grey region (bottom left) the model neuron did not fire. (B) The standard deviation of the latency (across trials) of the first thalamocortical spike relative the movement-related decrease distinguished rebound from disinhibition transmission modes. For the rebound mode (i.e. yellow area in A) the standard deviation was almost always the lowest, and the regime in which rebound and disinhibition coexisted the standard deviation was markedly higher. White contour line shows the boundaries of the yellow area in panel (A), where the transmission was exclusively mediated by rebound spiking. (C-E) Sample firing rate profiles and corresponding raster plots show the activity of the thalamocortical neuron in different parts of the parameter regime (as indicated by the corresponding letters in A) with rebound spiking only (C), coexistence of rebound and disinhibition (D-E) and disinhibition only (F).

We characterised the nigrothalamic transmission mode (see Materials and Methods) according to the proportion of trials with rebound spikes for a range of inhibitory and excitatory inputs strengths (Figure 6A). Motor signals were transmitted via rebound spikes even in the presence of weak excitatory inputs (*g*_*CX*→*TC*_ ≤ 1.5 *nS/µm*^2^; Figure 6A). Interestingly, the transition from rebound to disinhibition mode was not abrupt, but there was a region where disinhibition and rebound spikes coexisted (Figure 6B). In these overlapping regions rebound spiking seemed to be the dominant firing pattern with a strong, transient firing rate increase in response to the movement-related decrease, a phenomenon which was already observed in anesthetised songbirds (Kojima and Doupe, 2009; Figure 6D, E; see also Discussion). We also examined the effects of varying the firing rate of the excitatory inputs (200, 500, and 1000 Hz). While the rebound and disinhibition spiking mode still overlapped, the corresponding parameter region was shifted towards lower excitatory conductances. For moderate excitatory input firing rates (100 and 200 Hz), rebound spiking occurred also in regions in which the model neuron was spontaneously active (Figure 6E). This overlap was present for spontaneous activity up to 3 Hz in line with the average spontaneous firing of motor thalamus neurons in rats during open-field behavior (Bosch-Bouju et al., 2014). However, for higher spontaneous activity (>7 Hz) rebound spiking vanished (Figure 6F). We conclude that the model neuron can transmit motor signals in the rebound mode also in the presence of excitatory inputs.

We also characterised the transmission precision for different transmission modes by computing the standard deviation of the timing of the first spike after the movement-related decrease across trials (Figure 6B). For the rebound transmission mode, the transmission precision was maximal (i.e. minimal timing standard deviation), but as the proportion of trials with disinhibition mode increased, the transmission precision decreased. In the weak inhibition and excitation regime, where rebound and disinhibition modes coexisted and the baseline firing rate of the model neuron was low (< 7 Hz), the precision was smallest. This is important because the spiking variability can be characterised in electrophysiological recordings and may thus provide an indication of the transmission mode *in vivo*.

In summary, our computational model points to new functional roles for uncorrelated basal ganglia output in the high-fidelity transmission of motor signals. We characterised conditions and parameter regimes under which rebound spikes can occur in the thalamus as a response to movement-related decreases in firing rate of basal ganglia output neurons. In our model neuron, rebound spiking requires that most input from basal ganglia neurons exhibits movement-related pauses or synchronous baseline activity, and that inhibitory inputs are strong relative to the excitatory inputs. Therefore, our model is in line with previous studies arguing that the conditions for rebound spiking are primarily given in pathological situations. However, our model also points to the possibility that rebound spikes could occur in healthy animals, but only under rather specific conditions with respect to the connectivity and inputs, in a small subset of thalamic neurons.

## Discussion

We used computational modelling to study the impact of spike train correlations in the basal ganglia output on the transmission of motor signals. Based on previous studies we focused our description on movement-related pauses in SNr activity (Hikosaka and Wurtz, 1983; Schultz, 1986; Leblois et al., 2007; Schmidt et al., 2013) and examined their effect on motor thalamus in the rebound transmission mode. However, as also neurons in e.g. the superior colliculus can respond with a rebound spike after prolonged hyperpolarisation (Saito and Isa, 1999), our modelling results might apply more generally. Furthermore, while previous studies identified the important role of excitation in determining regimes in which rebound spikes can occur (Goldberg et al., 2013; Edgerton and Jaeger, 2014), our model produced rebound spikes in a wider parameter regime, also in the presence of excitation (Figure 6). In addition, rebound spiking overlapped with the disinhibition transmission mode, indicating that the different transmission modes might not always be clearly separable. In our model, the impaired nigrothalamic transmission of motor signals for correlated inputs also indicates a potential functional role of uncorrelated activity in basal ganglia output regions, possibly as a result of active decorrelation (Wilson, 2013).

While we focussed here on the conditions and properties of rebound spiking, our model did not support that rebound spiking is the dominant nigrothalamic transmission mode in healthy animals. In line with previous studies, we found that rebound spiking primarily occurs for correlated activity (mimicking pathological conditions). Furthermore, rebound spiking depended on three main parameters of the nigrothalamic connections: the strength of inhibition, the degree of convergence, and the number of rate-decreasing SNr neurons. Currently, the values of these parameters are unknown, but experimental studies provide rough estimates, and our computational approach allowed us to determine the range of parameter values under which rebound spikes can occur. In addition, basal ganglia output neurons have heterogeneous firing patterns during motor tasks (Hikosaka and Wurtz, 1983; Schultz, 1986; Leblois et al., 2007; Schmidt et al., 2013; Schwab et al., 2020). Therefore, it seems plausible that also the transmission mode may vary across neurons and over time. Accordingly, rebound spiking may only occur under specific circumstances in motor thalamus and involve only a subset of neurons. Similarly, our modelling results indicated that there may be specific conditions in which rebound spiking could also occur in healthy animals. For example, synchronous movement-related pauses in many SNr neurons might constitute a transient signal that could lead to rebound spiking in a subset of neurons in the motor thalamus. However, this would require strong and specific connections from SNr to thalamus so that SNr neurons with the same firing pattern converge onto the same thalamocortical neuron. As these rebound spikes would only form a small subset of the total number of thalamic spikes, it might be difficult to detect them in extracellular recordings (Schwab et al., 2020).

Our modelling results also indicated that rebound spiking depends on the baseline firing rate of motor thalamus neurons. Notably, neurons with a low baseline firing rate may be more likely to transmit motor signals via rebound spikes (Fig. 6A). From experimental studies we know that neurons in the motor thalamus have diverse baseline firing rates (Guo et al., 2017; Gaidica et al., 2018). Therefore, we would predict that the subset of neurons with low baseline firing rates are more likely to exhibit rebound spikes, when receiving synchronous inputs. In addition, the nigrothalamic connection strength was an important parameter in our simulations, with stronger connections favouring rebound spike generation (Fig. 6A; see also Kuramoto et al., 2011). Therefore, any change in nigrothalamic connection strength, e.g. during motor learning, can also affect the propensity of the circuit to generate rebound spikes.

Experimental studies have shown that rebound activity often involves bursts consisting of several spikes (Magnin et al., 2000; Bosch-Bouju et al., 2014; Edgerton and Jaeger, 2014). In contrast, our model here usually responded only with a single rebound spike. In our model this had the advantage to simplify our quantification of the transmission quality, which would require additional measures to classify spikes within a rebound burst. Importantly, in a model variation with rebound bursts the overall effect of correlations on the transmission quality would stay the same (Supplemental Figure 3). While rebound bursts in motor thalamus might play a role for transmitting motor signals further downstream, this is beyond the scope of this paper.

### Functional role of active decorrelation in the basal ganglia

One prominent feature of neural activity in the healthy basal ganglia is the absence of spike correlations (Bar-Gad et al., 2003). This might be due to the autonomous pacemaking activity of neurons in globus pallidus externa/interna (GPe/GPi), subthalamic nucleus (STN) and SNr, as well as other properties of the network such as heterogeneity of firing rates and connectivity that actively counteracts the synchronisation of activity (Wilson, 2013). While uncorrelated basal ganglia activity may maximise information transmission (Wilson, 2015), our simulations demonstrate that it further prevents the occurrence of random pauses in SNr/GPi activity that could drive thalamic rebound spikes. Thereby, uncorrelated basal ganglia output activity may ensure that rebound spikes in motor thalamus neurons do usually not occur in healthy animals, or only occur upon appropriate signals such as the movement-related decreases in basal ganglia output firing rate. In contrast, correlated basal ganglia output activity leads to rebound activity in motor thalamus also at baseline SNr activity, i.e. in absence of any motor signal. This decrease in the signal-to-noise ratio of motor signals may cause problems in motor control.

Evidence for the functional relevance of uncorrelated basal ganglia activity originates from the prominent observation that basal ganglia activity becomes correlated in Parkinson’s disease (Bergman et al., 1998; Nevado-Holgado et al., 2014). Therefore, our simulations with correlated basal ganglia output activity capture a key aspect of neural activity in Parkinson’s disease. Interestingly, our finding that basal ganglia correlations increase the rate of motor thalamus rebound spikes is in line with recent experimental findings. In dopamine-depleted mice with Parkinson-like motor symptoms, the rate of motor thalamus rebound spikes was also increased compared to healthy controls (Kim et al., 2017). Furthermore, an increased trial-to-trial variability of rebound spikes was found in dopamine-depleted mice, similar to our simulations (Figure 3).

Therefore, our results demonstrate the role of uncorrelated activity in the high-fidelity transmission of motor signals with low trial-to-trial variability from the basal ganglia to motor thalamus. Uncorrelated activity could be the result of active decorrelation in the basal ganglia (Wilson, 2013). In contrast, pathological correlations may lead to unreliable and noisy transmission of motor signals with high trial-to-trial variability, potentially contributing to motor symptoms in Parkinson’s disease.

### Role of rebound spikes for motor output

In our simulations we only examined the activity of a single thalamocortical neuron. However, for motor signals propagating further downstream, the coordination of activity among different thalamocortical neurons might be relevant. Due to the low trial-to-trial variability of the response latency of rebound spikes (Fig. 6B), in the model pauses in population SNr activity would lead to synchronous rebound spikes among thalamocortical neurons. In contrast, excitatory, Poisson inputs from cortex enhanced trial-to-trial variability (Fig. 6B) and thus would not lead to synchronous activity among thalamocortical neurons. Even though downstream regions cannot directly distinguish thalamic rebound spikes from excitation-driven spikes, they might read out synchronous activity that occurs primarily for rebound spikes. Thereby, only coordinated activity in different thalamocortical neurons may lead to movement initiation (Gaidica et al., 2018) or muscle contraction (Kim et al., 2017). This is in line with the experimental finding showing that, despite no significant difference in the peak or average firing rates of single unit recordings from intact and knockout neurons lacking T-type Ca^2+^ in the motor thalamus, multi-unit recordings from intact neurons reached a stronger peak firing rate earlier than the knockout neurons (Kim et al., 2017). This early activation of a greater proportion of intact neurons after the termination of the inhibition, which indicates a coordinated activity across neurons, was accompanied by a muscular response whereas no muscular response was observed in the knockout state (Kim et al., 2017). Therefore, rebound activity in an individual motor thalamus neuron may not lead to muscle contraction, but instead synchronous rebound spikes in several motor thalamus neurons may be required.

### Impact of sensory responses on the transmission of motor signals

SNr neurons that decrease their activity during movement also respond to salient sensory stimuli such as auditory “Go” stimuli cueing movement (Pan et al., 2013; Schmidt et al., 2013). One proposed functional role for this brief firing rate increase is to prevent impulsive or premature responses during movement preparation in SNr neurons (Schmidt et al., 2013). In addition, in our model we observed that, depending on the precise timing, inhibitory inputs mimicking sensory responses may also promote thalamocortical rebound spikes. This effect was present in the model when the sensory responses preceded the movement-related decrease by up to 40 ms (Figure 4).

In rats performing a stop-signal task the same SNr neurons that responded to the “Go” stimulus also responded to an auditory “Stop” signal, which prompted the cancellation of the upcoming movement (Schmidt et al., 2013). These responses were observed in trials, in which the rats correctly cancelled the movement, but not in trials where they failed to cancel the movement. These SNr responses to the “Stop” signal may delay movement initiation, allowing another slower process to completely cancel the planned movement (Mallet et al., 2016). In line with this “pause-then-cancel” model of stopping (Schmidt and Berke, 2017), we observed that the SNr sensory responses can also prevent rebound spikes when they occur close to the time of the motor signal. In our model this suppression effect was present up to 40 ms after the onset of the movement-related decrease in SNr activity (Figure 4). Thereby, our model provides a prediction for the temporal window of the functional contribution of sensory responses in SNr to behaviour. Importantly, sensory responses could either promote or suppress movements, depending on their relative timing to the motor signal, providing a highly flexible means to integrate sensory and motor signals in nigrothalamic circuits. However, due to the restricted parameter range in which our model generated rebound spikes, it is unclear whether the modulation of rebound spiking by sensory responses could also occur in healthy animals.

### Effects of deep brain stimulation

In our model correlated basal ganglia activity increased the number of rebound spikes in thalamocortical neurons. In particular, higher-order correlations lead to pauses in the SNr population activity promoting rebound spikes, while pairwise correlations alone did not affect the nigrothalamic transmission of motor signals (Figure 2B). This suggests that in Parkinson’s disease higher-order correlations are relevant for motor symptoms, which offers some insight into the potential mechanisms by which deep-brain stimulation (DBS) might alleviate some of the motor symptoms such as rigidity and tremor. DBS in the STN and GPi has complex and diverse effects on the firing rate of neurons in SNr/GPi (Bar-Gad et al., 2004; Zimnik et al., 2015) and thalamus (Muralidharan et al., 2017). According to our model strong increases in SNr and GPi firing rates observed after STN DBS (Hashimoto et al., 2003; Maurice et al., 2003), would decrease the duration of the spontaneous pauses in the population activity (Figure 3C). Thereby, even for correlated SNr activity, the duration of the pauses would not be long enough to allow the generation of a rebound spike in the thalamocortical neuron. This conclusion also holds when a subset of neurons in SNr and GPi decrease their firing rate during STN DBS (Hahn et al., 2008; Humphries and Gurney, 2012). The decrease in the firing rate would decrease the degree of correlation by eliminating or displacing the synchronous spike times and therefore weaken the inhibition preceding the pauses that could have potentially evoked rebound spikes.

## Acknowledgements

This work was supported by Erasmus Mundus joint PhD program (EuroSPIN), the BrainLinks-BrainTools Cluster of Excellence funded by the German Research Foundation (DFG, grant number: EXC 1086), the EU H2020 Programme as part of the Human Brain Project (HBP-SGA1, 720270; HBP-SGA2, 785907), and the University of Sheffield. We also acknowledge support by the state of Baden-Wuerttemberg through bwHPC and the German Research Foundation (DFG) through grant no INST 39/963-1 FUGG. We would like to thank David Bilkey, Alejandro Jimenez, Lars Hunger, Amin Mirzaei, and Genela Morris for helpful discussions.

## Author Contributions

Mohammadreza Mohagheghi Nejad and Robert Schmidt designed the research. Robert Schmidt supervised the work. Mohammadreza Mohagheghi Nejad performed the simulations and analysed the data. Mohammadreza Mohagheghi Nejad, Stefan Rotter and Robert Schmidt interpreted the results and wrote the manuscript.

## Data Accessibility

We provided our simulation scripts (in “BasicModelSimulations” directory) including the scripts generating input spike trains (in “SpikeTrains” directory) accessible via a git repository https://github.com/mmohaghegh/NigrothalamicTransmission.git.

### Abbreviations

BIN: Binomially distributed
CX: Cortex
DBS: Deep Brain Stimulation
DLM: Dorsolateral nucleus of the anterior thalamus
EXP: Exponentially distributed
GABA: gamma-Aminobutyric acid
GPe: Globus Pallidus externa
GPi: Globus Pallidus interna
SNr: Substantia Nigra pars reticulata
STN: Subthalamic Nucleus
TC: Thalamocortical neuron
TQ: Transmission Quality

## Figure Legends

**Supplemental Figure 1.**
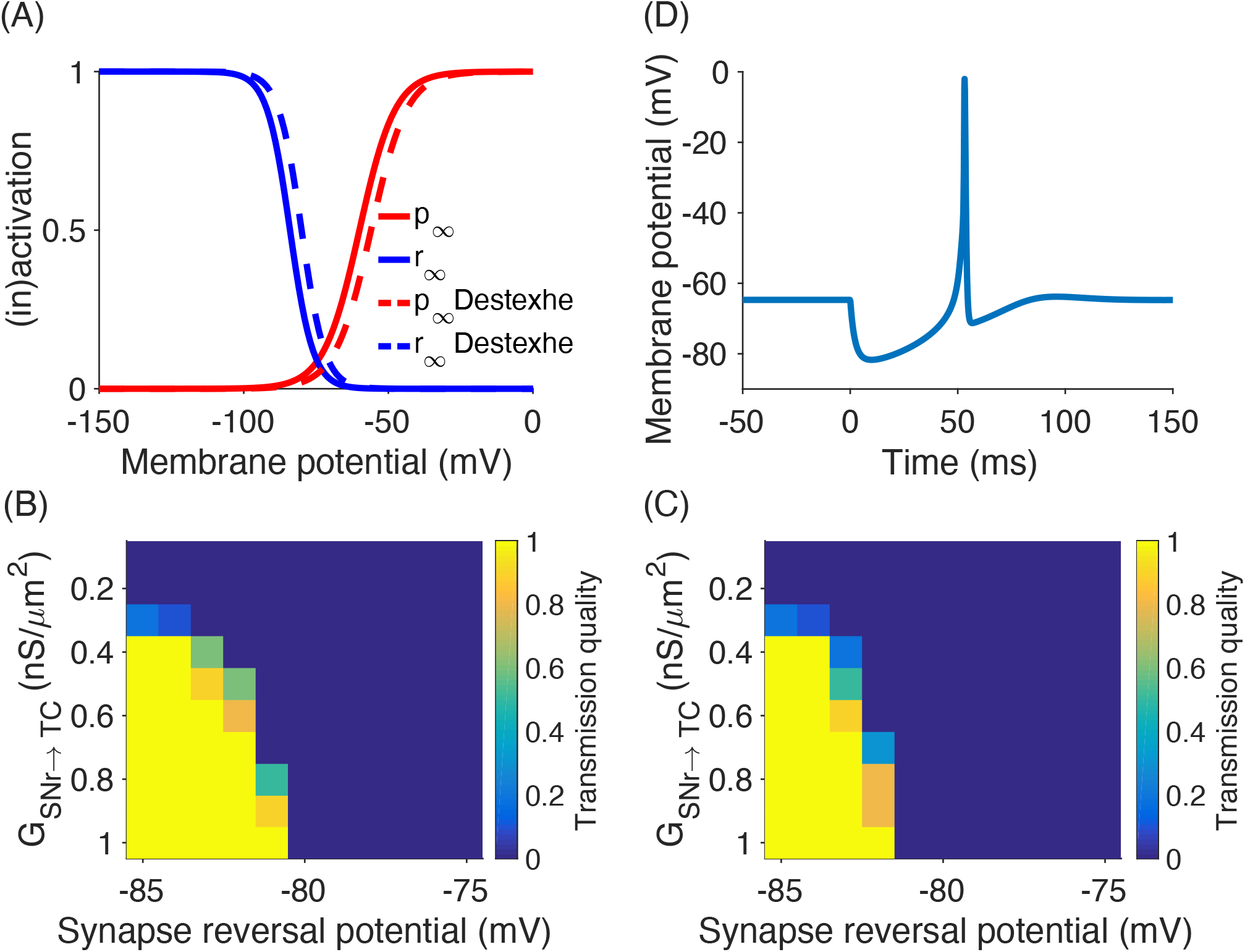
Robustness of the model response to inhibitory synapse parameters. (A) Comparison of gating variable equilibrium values as a function of the membrane potential between our model and the one from Destexhe et al. (1998). (B) Transmission quality of our model measured in response to 30 synaptic inputs that decrease their activity from 50Hz to 0 at the time of movement onset. Our model produces high transmission quality also beyond the default parameter for reversal potential (−85*mV*). (C) Adapting the settings from Destexhe et al. (1998) for gating variables equilibrium values for our model only slightly decreases the transmission quality in comparison to the default parameter settings in panel (B). (D) Exposing the model neuron to 30 synchronous spikes arriving at the synapse (GABA maximum conductance = 1*nS/µm*^2^) at time 0ms hyperpolarises the membrane potential from −64.7*mV* to −81.7*mV*. This hyperpolarisation is strong enough to evoke a post-inhibitory rebound spike in our model, which is very close to the hyperpolarisation (18*mV*) of thalamocortical neurons in motor thalamus in vitro (see Figure 5B in Edgerton and Jaeger (2014)).

**Supplemental Figure 2.**
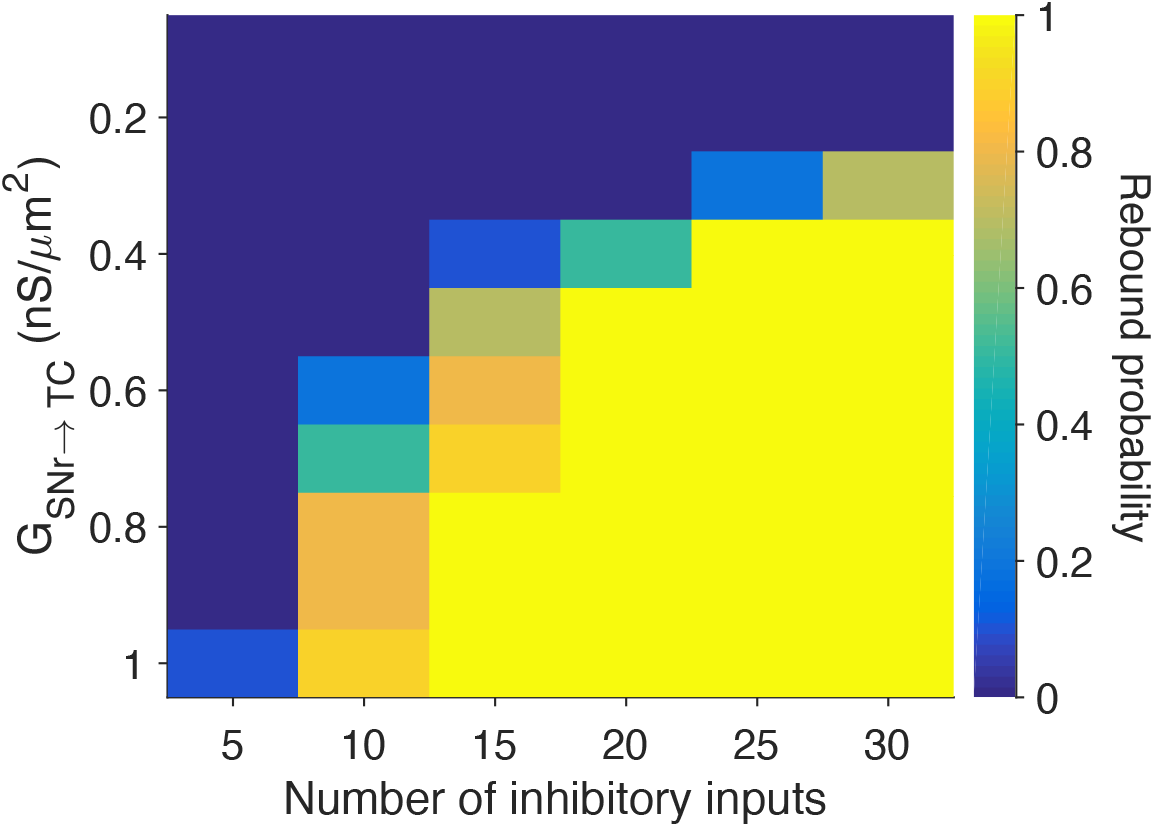
The effect of the number of inhibitory inputs on the probability of rebound spike generation. Reducing the number of inputs from 30 (default value used throughout this study) to 20 does not substantially change the probability of rebound spikes in the model. However, to generate rebound spikes with high probability for less than 20 inputs, the model requires stronger inhibition (higher *G*_*SNr→TC*_).

**Supplemental Figure 3.**
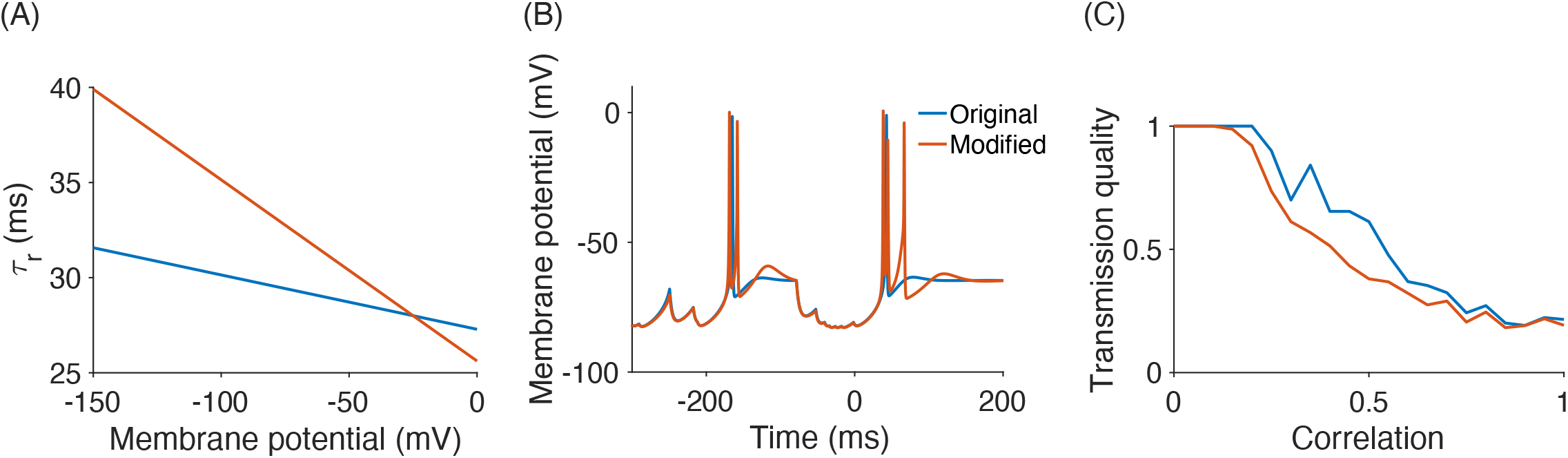
Modified model in which rebound spiking involves a burst of action potentials (original model in blue, modified model in red). (A) The modified model had a higher voltage-dependent time constant of the inactivation gating variable *τ*_*r*_ for most values for the model membrane potential. (B) Simulation with correlated inhibitory inputs firing at 50*Hz* and a movement-related decrease in the firing rates. The increase in the time constant in the modified model leads to rebound bursts (red line), as can be seen by the additional spikes compared to the original model (blue line). (C) The modified model preserved the overall decrease in the transmission quality as a function of correlation, with a slightly lower transmission quality for intermediate correlations in the modified model.

## References

Albin RL, Young AB, Penney JB (1989) The functional anatomy of basal ganglia disorders. Trends in Neurosciences 12:366–375.

Alexander GE, Crutcher MD (1990a) Functional architecture of basal ganglia circuits: neural substrates of parallel processing. Trends in Neurosciences 13:266–271.

Alexander GE, Crutcher MD (1990b) Neural representations of the target (goal) of visually guided arm movements in three motor areas of the monkey. Journal of Neurophysiology 64:164–178.

Avila I, Parr-Brownlie LC, Brazhnik E, Castañeda E, Bergstrom DA, Walters JR (2010) Beta frequency synchronization in basal ganglia output during rest and walk in a hemiparkinsonian rat. Experimental Neurology 221:307–319.

Bar-Gad I, Elias S, Vaadia E, Bergman H (2004) Complex locking rather than complete cessation of neuronal activity in the globus pallidus of a 1-methyl-4-phenyl-1, 2, 3, 6-tetrahydropyridine-treated primate in response to pallidal microstimulation. Journal of Neuroscience 24:7410–7419.

Bar-Gad I, Heimer G, Ritov Y, Bergman H (2003) Functional correlations between neighboring neurons in the primate globus pallidus are weak or nonexistent. Journal of Neuroscience 23:4012–4016.

Bergman H, Feingold A, Nini A, Raz A, Slovin H, Abeles M, Vaadia E (1998) Physiological aspects of information processing in the basal ganglia of normal and parkinsonian primates. Trends in Neurosciences 21:32–38.

Bodor ÁL, Giber K, Rovó Z, Ulbert I, Acsády L (2008) Structural correlates of efficient gabaergic transmission in the basal ganglia–thalamus pathway. Journal of Neuroscience 28:3090–3102.

Bosch-Bouju C, Hyland BI, Parr-Brownlie LC (2013) Motor thalamus integration of cortical, cerebellar and basal ganglia information: implications for normal and parkinsonian conditions. Frontiers in Computational Neuroscience 7:163.

Bosch-Bouju C, Smither RA, Hyland BI, Parr-Brownlie LC (2014) Reduced reach-related modulation of motor thalamus neural activity in a rat model of parkinson’s disease. Journal of Neuroscience 34:15836–15850.

Brown P, Oliviero A, Mazzone P, Insola A, Tonali P, Di Lazzaro V (2001) Dopamine dependency of oscillations between subthalamic nucleus and pallidum in parkinson’s disease. Journal of Neuroscience 21:1033–1038.

Bujan AF, Aertsen A, Kumar A (2015) Role of input correlations in shaping the variability and noise correlations of evoked activity in the neocortex. Journal of Neuroscience 35:8611–8625.

Daw ND, Niv Y, Dayan P (2005) Uncertainty-based competition between prefrontal and dorsolateral striatal systems for behavioral control. Nature Neuroscience 8:1704.

Deniau J, Chevalier G (1985) Disinhibition as a basic process in the expression of striatal functions. ii. the striato-nigral influence on thalamocortical cells of the ventromedial thalamic nucleus. Brain Research 334:227–233.

Destexhe A, Contreras D, Steriade M (1998) Mechanisms underlying the synchronizing action of corticothalamic feedback through inhibition of thalamic relay cells. Journal of Neurophysiology 79:999–1016.

Edgerton JR, Jaeger D (2014) Optogenetic activation of nigral inhibitory inputs to motor thalamus in the mouse reveals classic inhibition with little potential for rebound activation. Frontiers in Cellular Neuroscience 8:36.

Ermentrout GB, Terman DH (2010) Mathematical foundations of neuroscience, Vol. 35 Springer Science & Business Media.

Gaidica M, Hurst A, Cyr C, Leventhal DK (2018) Distinct populations of motor thalamic neurons encode action initiation, action selection, and movement vigor. Journal of Neuroscience.

Gerstner W, Kistler WM (2002) Spiking neuron models: Single neurons, populations, plasticity Cambridge University Press.

Goldberg JH, Farries MA, Fee MS (2012) Integration of cortical and pallidal inputs in the basal ganglia-recipient thalamus of singing birds. Journal of Neurophysiology 108:1403–1429.

Goldberg JH, Farries MA, Fee MS (2013) Basal ganglia output to the thalamus: still a paradox. Trends in Neurosciences 36:695–705.

Goldberg JH, Fee MS (2012) A cortical motor nucleus drives the basal ganglia-recipient thalamus in singing birds. Nature Neuroscience 15:620–627.

Guo Y, Rubin JE, McIntyre CC, Vitek JL, Terman D (2008) Thalamocortical relay fidelity varies across subthalamic nucleus deep brain stimulation protocols in a data-driven computational model. Journal of Neurophysiology 99:1477–1492.

Guo ZV, Inagaki HK, Daie K, Druckmann S, Gerfen CR, Svoboda K (2017) Maintenance of persistent activity in a frontal thalamocortical loop. Nature 545:181–186.

Haber SN, Calzavara R (2009) The cortico-basal ganglia integrative network: the role of the thalamus. Brain Research Bulletin 78:69–74.

Hahn PJ, Russo GS, Hashimoto T, Miocinovic S, Xu W, McIntyre CC, Vitek JL (2008) Pallidal burst activity during therapeutic deep brain stimulation. Experimental Neurology 211:243–251.

Hashimoto T, Elder CM, Okun MS, Patrick SK, Vitek JL (2003) Stimulation of the subthalamic nucleus changes the firing pattern of pallidal neurons. Journal of Neuroscience 23:1916–1923.

Herd MB, Brown AR, Lambert JJ, Belelli D (2013) Extrasynaptic gabaa receptors couple presynaptic activity to postsynaptic inhibition in the somatosensory thalamus. Journal of Neuroscience 33:14850–14868.

Hikosaka O, Takikawa Y, Kawagoe R (2000) Role of the basal ganglia in the control of purposive saccadic eye movements. Physiological Reviews 80:953–978.

Hikosaka O, Wurtz RH (1983) Visual and oculomotor functions of monkey substantia nigra pars reticulata. iv. relation of substantia nigra to superior colliculus. Journal of Neurophysiology 49:1285–1301.

Huguenard J, Prince D (1994) Intrathalamic rhythmicity studied in vitro: nominal t-current modulation causes robust antioscillatory effects. Journal of Neuroscience 14:5485–5502.

Humphries MD, Gurney K (2012) Network effects of subthalamic deep brain stimulation drive a unique mixture of responses in basal ganglia output. European Journal of Neuroscience 36:2240–2251.

Kase D, Uta D, Ishihara H, Imoto K (2015) Inhibitory synaptic transmission from the substantia nigra pars reticulata to the ventral medial thalamus in mice. Neuroscience Research 97:26–35.

Kim J, Kim Y, Nakajima R, Shin A, Jeong M, Park AH, Jeong Y, Jo S, Yang S, Park H et al. (2017) Inhibitory basal ganglia inputs induce excitatory motor signals in the thalamus. Neuron 95:1181–1196.

Kojima S, Doupe AJ (2009) Activity propagation in an avian basal ganglia-thalamocortical circuit essential for vocal learning. Journal of Neuroscience 29:4782–4793.

Kravitz AV, Freeze BS, Parker PR, Kay K, Thwin MT, Deisseroth K, Kreitzer AC (2010) Regulation of parkinsonian motor behaviours by optogenetic control of basal ganglia circuitry. Nature 466:622–626.

Kuhn A, Aertsen A, Rotter S (2003) Higher-order statistics of input ensembles and the response of simple model neurons. Neural Computation 15:67–101.

Kumar A, Cardanobile S, Rotter S, Aertsen A (2011) The role of inhibition in generating and controlling parkinson’s disease oscillations in the basal ganglia. Frontiers in Systems Neuroscience 5:86.

Kuramoto E, Fujiyama F, Nakamura KC, Tanaka Y, Hioki H, Kaneko T (2011) Complementary distribution of glutamatergic cerebellar and gabaergic basal ganglia afferents to the rat motor thalamic nuclei. European Journal of Neuroscience 33:95–109.

Laudes T, Meis S, Munsch T, Lessmann V (2012) Impaired transmission at corticothalamic excitatory inputs and intrathalamic gabaergic synapses in the ventrobasal thalamus of heterozygous bdnf knockout mice. Neuroscience 222:215–227.

Leblois A, Bodor ÁL, Person AL, Perkel DJ (2009) Millisecond timescale disinhibition mediates fast information transmission through an avian basal ganglia loop. Journal of Neuroscience 29:15420–15433.

Leblois A, Meissner W, Bezard E, Bioulac B, Gross CE, Boraud T (2006) Temporal and spatial alterations in gpi neuronal encoding might contribute to slow down movement in parkinsonian monkeys. European Journal of Neuroscience 24:1201–1208.

Leblois A, Meissner W, Bioulac B, Gross CE, Hansel D, Boraud T (2007) Late emergence of synchronized oscillatory activity in the pallidum during progressive parkinsonism. European Journal of Neuroscience 26:1701–1713.

Lindahl M, Kotaleski JH (2016) Untangling basal ganglia network dynamics and function–role of dopamine depletion and inhibition investigated in a spiking network model. Eneuro 3:ENEURO.0156–16.2016.

Llinás R, Jahnsen H (1982) Electrophysiology of mammalian thalamic neurones in vitro. Nature 297:406.

Magnin M, Morel A, Jeanmonod D (2000) Single-unit analysis of the pallidum, thalamus and subthalamic nucleus in parkinsonian patients. Neuroscience 96:549–564.

Mallet N, Schmidt R, Leventhal D, Chen F, Amer N, Boraud T, Berke JD (2016) Arkypallidal cells send a stop signal to striatum. Neuron 89:308–316.

Maurice N, Thierry AM, Glowinski J, Deniau JM (2003) Spontaneous and evoked activity of substantia nigra pars reticulata neurons during high-frequency stimulation of the subthalamic nucleus. Journal of Neuroscience 23:9929–9936.

Mirzaei A, Kumar A, Leventhal D, Mallet N, Aertsen A, Berke J, Schmidt R (2017) Sensorimotor processing in the basal ganglia leads to transient beta oscillations during behavior. Journal of Neuroscience 37:1289–17.

Muralidharan A, Zhang J, Ghosh D, Johnson MD, Baker KB, Vitek JL (2017) Modulation of neuronal activity in the motor thalamus during gpi-dbs in the mptp nonhuman primate model of parkinson’s disease. Brain Stimulation 10:126–138.

Nevado-Holgado AJ, Mallet N, Magill PJ, Bogacz R (2014) Effective connectivity of the subthalamic nucleus–globus pallidus network during parkinsonian oscillations. The Journal of Physiology 592:1429–1455.

Pan WX, Brown J, Dudman JT (2013) Neural signals of extinction in the inhibitory microcircuit of the ventral midbrain. Nature Neuroscience 16:71.

Person AL, Perkel DJ (2005) Unitary ipsps drive precise thalamic spiking in a circuit required for learning. Neuron 46:129–140.

Person AL, Perkel DJ (2007) Pallidal neuron activity increases during sensory relay through thalamus in a songbird circuit essential for learning. Journal of Neuroscience 27:8687–8698.

Redgrave P, Prescott TJ, Gurney K (1999) The basal ganglia: a vertebrate solution to the selection problem? Neuroscience 89:1009–1023.

Redgrave P, Rodriguez M, Smith Y, Rodriguez-Oroz MC, Lehericy S, Bergman H, Agid Y, DeLong MR, Obeso JA (2010) Goal-directed and habitual control in the basal ganglia: implications for parkinson’s disease. Nature Reviews Neuroscience 11:760.

Reitsma P, Doiron B, Rubin JE (2011) Correlation transfer from basal ganglia to thalamus in parkinson’s disease. Frontiers in Computational Neuroscience 5:58.

Rinzel J (1985a) Excitation dynamics: insights from simplified membrane models In Fed. Proc, Vol. 44, pp. 2944–2946.

Rinzel J (1985b) Excitation dynamics: insights from simplified membrane models In Fed. Proc, Vol. 44, pp. 2944–2946.

Rubin JE, Terman D (2004) High frequency stimulation of the subthalamic nucleus eliminates pathological thalamic rhythmicity in a computational model. Journal of Computational Neuroscience 16:211–235.

Saito Y, Isa T (1999) Electrophysiological and morphological properties of neurons in the rat superior colliculus. i. neurons in the intermediate layer. Journal of Neurophysiology 82:754–767.

Schmidt R, Berke JD (2017) A pause-then-cancel model of stopping: evidence from basal ganglia neurophysiology. Phil. Trans. R. Soc. B 372:20160202.

Schmidt R, Leventhal DK, Mallet N, Chen F, Berke JD (2013) Canceling actions involves a race between basal ganglia pathways. Nature Neuroscience 16:1118.

Schultz W (1986) Activity of pars reticulata neurons of monkey substantia nigra in relation to motor, sensory, and complex events. Journal of Neurophysiology 55:660–677.

Schwab BC, Kase D, Zimnik A, Rosenbaum R, Codianni MG, Rubin JE, Turner RS (2020) Neural activity during a simple reaching task in macaques is counter to gating and rebound in basal ganglia–thalamic communication. PLOS Biology 18:1–38.

Staude B, Grün S, Rotter S (2010) Higher-order correlations and cumulants In Analysis of parallel spike trains, pp. 253–280. Springer.

Ulrich D, Huguenard JR (1997) Nucleus-specific chloride homeostasis in rat thalamus. Journal of Neuroscience 17:2348–2354.

Wichmann T, DeLong MR (1996) Functional and pathophysiological models of the basal ganglia. Current Opinion in Neurobiology 6:751–758.

Wilson CJ (2013) Active decorrelation in the basal ganglia. Neuroscience 250:467–482.

Wilson CJ (2015) Oscillators and oscillations in the basal ganglia. The Neuroscientist 21:530–539.

Yin HH, Knowlton BJ (2006) The role of the basal ganglia in habit formation. Nature Reviews Neuroscience 7:464.

Zimnik AJ, Nora GJ, Desmurget M, Turner RS (2015) Movement-related discharge in the macaque globus pallidus during high-frequency stimulation of the subthalamic nucleus. Journal of Neuroscience 35:3978–3989.

